# Interleukin-6 elevates thrombosis via pro-coagulant phospholipids from platelet 12-lipoxygenase in rheumatoid arthritis

**DOI:** 10.1101/2025.03.26.645440

**Authors:** Daniela O Costa, Stuart T.O Hughes, Robert H. Jenkins, Ana Cardus Figueras, Majd B Protty, Victoria J Tyrrell, Ali A Hajeyah, Gareth W Jones, James J Burston, Beth Morgan, Federica Monaco, David Hill, Aisling S Morrin, Carol Guy, Alice Bacon, Martin Giera, Rene E M Toes, P Vince Jenkins, Peter W Collins, Ernest Choy, Simon A. Jones, Valerie B O’Donnell

## Abstract

**Background:** Rheumatoid arthritis (RA) is associated with significantly higher thrombotic risk, which is not yet mechanistically understood. Here, the role of pro-coagulant membranes of platelets and blood cells in driving thrombosis, and their regulation by inflammation was determined using human cohorts and genetically-modified mice.

**Methods:** Antigen-induced arthritis (AIA) was induced in *WT, Il27ra^−/−^, Il6ra*^−/−^, *Alox12^−/−^* and *Alox15*^−/−^ mice. Coagulation and inflammatory markers were measured in plasma. Lipidomics was performed on blood cells and synovium analyzing pro-coagulant enzymatically-oxidized phospholipids (eoxPL) and oxylipins. Two human RA patient cohorts were characterized for eoxPL generation in blood cells, and chronic immune response to eoxPL in vivo.

**Results:** AIA induction significantly elevated plasma thrombin-antithrombin (TAT) complexes, serum amyloid A (SAA), and eoxPL in blood cells and platelets. Elevations in TATs, SAA and eoxPL were suppressed by genetic deletion of IL-6Ra, while platelet *Alox12* deletion prevented TAT and eoxPL increases. This indicates a direct role for IL-6 in elevating thrombosis via upregulation of platelet eoxPL. In contrast, leukocyte *Alox15* deletion did not impact TATs or eoxPL. Deletion of either LOX isoform worsened AIA joint pathology. Synovial tissue demonstrated raised eoxPL, but exclusively from *Alox15*, indicating leukocyte origin. Thus, both LOX isoforms contribute to AIA, but through different mechanisms. In human RA, platelet counts, and plasma TATs were elevated, and plasma had significantly elevated IgG against eoxPL, indicating patients experience chronic exposure to the lipids *in vivo*.

**Conclusions:** Platelet-derived pro-coagulant eoxPL are elevated in human and murine arthritis along with higher coagulation markers. In mice, this was mediated by the IL-6/*Alox12* axis and directly responsible for the higher thrombotic risk. IL-6 plays a central role in driving platelet activation in RA, with the pro-coagulant lipid membrane representing a novel target. Reducing inflammation using DMARDs, particularly targeting IL-6 may reduce platelet pro-coagulant activity and thrombosis risk in RA.

## Introduction

Rheumatoid arthritis (RA) is associated with increased thrombotic risk, evidenced by 30-50% increased incidence of cardiovascular disease and stroke^1–4^. Coagulation is driven by circulating plasma factors which require an electronegative membrane, provided by platelets and white blood cells (WBC), following agonist activation. This comprises phospholipids (PL), namely phosphatidylserine (PS), phosphatidylethanolamine (PE) and enzymatically-oxidized phospholipids (eoxPL). These are either rapidly externalized (PS, PE)^5,6^ or generated de novo (eoxPL) following agonist activation^7–9^. EoxPL are formed via esterification of oxylipins generated by either lipoxygenases (LOX) or cyclooxygenases (COX) or direct oxygenation of PL or lysoPL^10,11^.

Humans with anti-phospholipid syndrome, an autoimmune disease with elevated thrombotic risk, display higher levels of circulating eoxPL^12^. This led us to ask whether the PL membrane is altered in other immune-mediated inflammatory diseases (IMIDs) such as RA and directly contributes to thrombosis. Limited studies on coagulation in RA have been undertaken. Several prothrombotic markers have been described as altered in patients^13^. Elevated thrombin activation, reflected by increased thrombin-antithrombin (TAT) complexes^14^ and D-dimers was described in RA plasma^15^. Tissue factor (TF) positively correlates with plasma C-Reactive Protein (CRP) and leukocyte counts, suggesting TF may play a role, and that thrombosis is associated with inflammation^14^. However, prothrombin and thrombin times are normal in RA, indicating that factor levels are not impacted^15^. This raises questions relating to how coagulation is regulated in RA and whether the pro-thrombotic membrane is involved.

Few studies have examined blood clotting in arthritis models, and no mechanistic research has been done. One study showed that on day 3 (early disease), plasma TF activity is elevated in murine antigen-induced arthritis (AIA), while TATs are elevated in collagen-induced arthritis (CIA)^16^. To address this information gap, studies in human cohorts and mouse models were undertaken here, focusing on the role of the pro-coagulant eoxPL membrane. In patients with RA, the levels and cellular origin of blood cell eoxPL were determined, while in serum, IgG-dependent immunoreactivity to eoxPL was characterized. Plasma from mice with AIA was analyzed for coagulation and fibrinolysis, and the role of inflammation driving production and action of eoxPL determined using genetically-modified mouse strains. Our study demonstrates that eoxPL from platelets drive the coagulation phenotype in arthritis, in an IL-6 dependent manner, representing a new therapeutic target for lowering thrombotic risk associated with chronic inflammatory disease.

## Methods

### Animal experiments

*Il27ra^−/−^* ^17^, *Il6ra^−/−^* ^18^, *Alox15^−/−^* and *Alox12^−/−^* ^19^ mice on C57BL/6 background were bred under specific pathogen-free conditions. Experiments were conducted in accordance with the United Kingdom Home Office Animals (Scientific Procedures) Act 1986, under project licenses PE8BCF782 and PC0174E40. AIA was induced (male, 9-12 weeks) as described ^20^, with full detail in Supplementary Methods. Whole joints were obtained and processed as outlined in Supplementary Methods for histology. Synovial tissue was dissected from the joint cavity and lipid extraction performed as in Supplementary Methods. Mouse blood collection and processing were performed as described^21^, with full detail in Supplementary Methods. Lipid extraction of whole blood cells, acid hydrolysis and LC-MS/MS analysis were conducted as in Supplementary Methods.

### Human studies

Patient studies were performed under three projects as outlined in Supplementary Methods. These were “Cardiff Regional Experimental Arthritis Treatment and Evaluation Centre”, approved by the Ethics Committee for Wales (Reference No 12/WA/0045), “Early Arthritis Cohort Biobank” (Leiden University Medical Center), and “Study of the lipidomic profile of blood clots from healthy volunteers” approved by School of Medicine Ethics Review Committee, Cardiff University (REC/SREC reference No 16/02, study 10). All followed the principles of the Declaration of Helsinki, with informed consent and full ethical approval. Methods for isolation of blood cells and platelets, activation, lipid extraction and analysis, as well as coagulation parameters are provided in Supplementary Methods.

## Results

### TATs and SAA are increased by IL-6 in murine arthritis

Coagulation parameters were determined in plasma on days 3 and 10 reflecting early and established arthritis. To map onto human phenotypes, the model was induced in strains which develop either myeloid-rich disease, driven by macrophages (WT), lymphoid-rich disease, with an elevated adaptive immune response (*Il27ra*^−/−^), or a fibroblast-rich phenotype, with reduced immune inflammation (*Il6ra^−/−^*)^22–26^. Immunization without arthritis induction, did not impact TATs, d-dimers, prothrombin time (PT) or C-reactive protein (CRP) (Supplementary Figure 1). Arthritis was confirmed by joint swelling in all 3 mouse strains following induction (Supplementary Figure 2 A). Platelet counts did not vary between strains, or during AIA (Supplementary Figure 2 B-C). TATs significantly increased (day 10), in both WT and *Il27ra^−/−^*, but not in *Il6ra^−/−^* mice (Figure 1 A). This indicates that coagulation is stimulated by IL-6-driven pathways in AIA. However, D-dimers or PT were not impacted at either timepoint (Supplementary Figure 3 A,B). Overall, while activation of coagulation occurred, there was no increase in fibrin degradation or consumptive coagulopathy. This is consistent with elevated thrombotic risk and similar to human RA, where TATs are elevated, but no alterations to PT or partial thromboplastin times are observed^15^.

**Figure 1.**
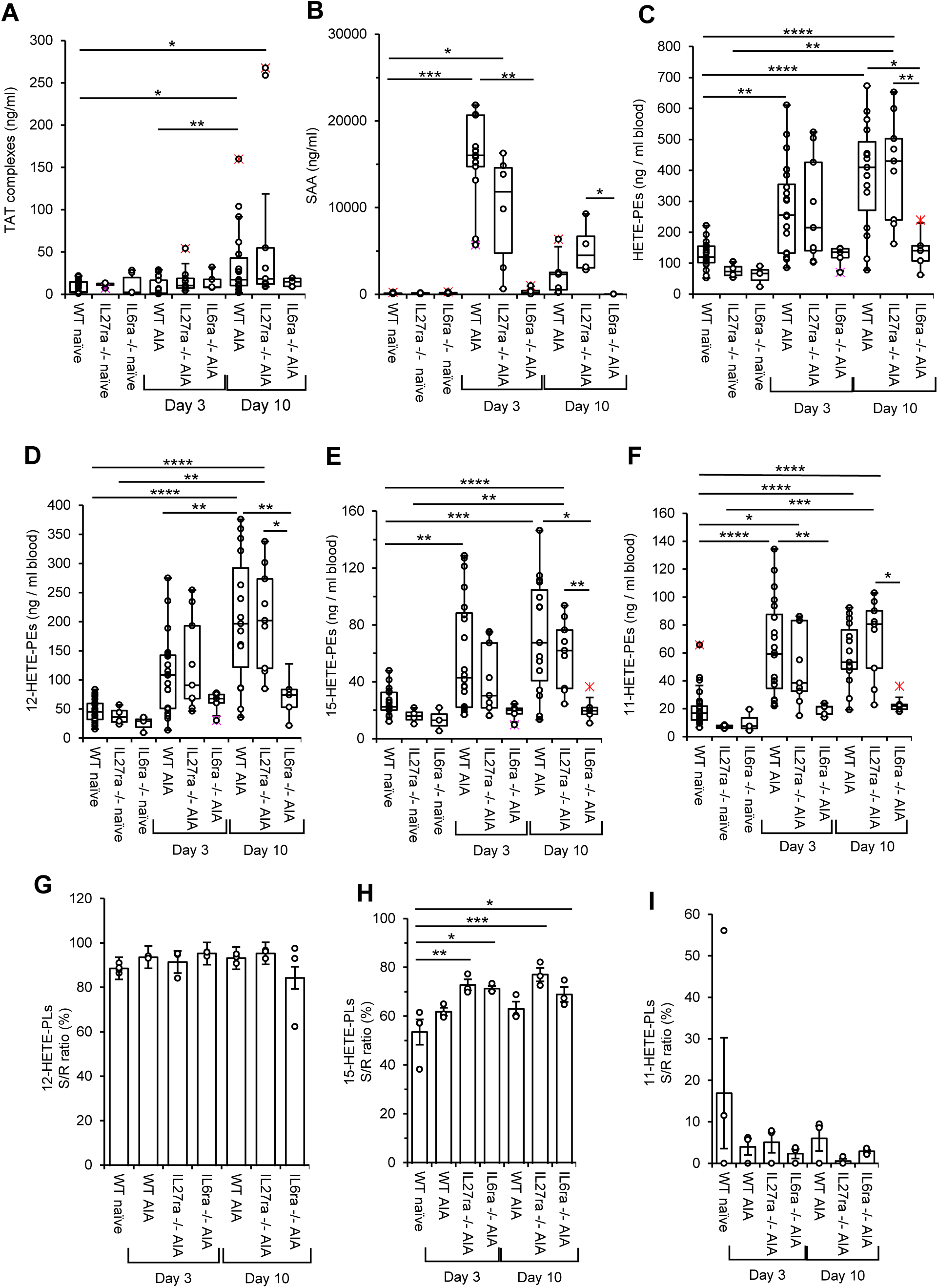
Murine arthritis increases TATs and platelet eoxPL in WT and *Il27ra*^−/−^ mice, but not *IL6ra*^−/−^. AIA was induced in 8-12 week old *WT, Il27ra*^−/−^ and *Il6ra*^−/−^ male mice as described in Methods, with whole blood collected on Days 3 and 10. *Panel A. TATs are elevated on Day 10 in AIA, but not in IL6ra^−/−^ mice.* TAT complexes were measured using ELISA. Plasma was collected on day 0 from WT naïve (n = 30), IL27ra^−/−^ naïve (n = 4) and IL6ra^−/−^ naïve (n = 4), as well as on days and 10 of AIA development from *WT (n = 19 and 18, respectively), IL27ra*^−/−^ (n= 9) and *IL6ra*^−/−^ (n= 5) mice. *Panel B. SAA is elevated in AIA, but not in IL6ra^−/−^ mice.* SAA was measured using ELISA. Plasma was collected on day 0 from WT naïve (n = 8), IL27ra^−/−^ naïve (n = 4) and IL6ra^−/−^ naïve (n = 4), as well as on days 3 and 10 of AIA development from *WT (n = 13 and 11,* respectively), *IL27ra*^−/−^ (*n = 6 and 5,* respectively) and *IL6ra*^−/−^ (n = 6 and 4, respectively) mice. *Panel C-F. Total sum of HETE-PL, along with 12-, 15-, and 11-HETE-PEs were significantly elevated in AIA, but not in IL6ra^−/−^ mice.* HETE-PL were quantified as outlined in Methods using LC/MS/MS. Whole blood was collected at day 0 from WT naïve (n = 23), *Il27ra^−/−^* naïve (n = 4) and *Il6ra^−/−^* naïve (n = 3), as well as, on days 3 and 10 of AIA development *WT* (n = 20 and 15, respectively), *Il27ra*^−/−^ (n = 9 for both days) and *Il6ra*^−/−^ (n = 5 for both days) male mice. Data is represented in box and whisker plot. *Panel G-F. Increased generation of 12-, 15- and 11-HETE-PL during AIA is enzymatic.* Chirality of eoxPL was determined in pooled lipid extracts as outlined in Methods using LC/MS/MS and expressed as S/R ratio (%) (n = 3) and represented by bar chart plot. Data were analyzed using Two-way ANOVA and Tukey’s multiple comparisons tests (*p<0.05, **p<0.01,*** p<0.001,**** p<0.0001).

To examine the role of inflammation, CRP and SAA were measured. SAA increased in WT and *Il27ra^−/−^* earlier than TATs, but not *Il6ra^−/−^* mice (Figure 1 B), consistent with known induction of SAA by IL-6^27^ and use of SAA as a biomarker of venous thromboembolism in humans^28^. Therefore, elevated TATs in WT and *Il27ra^−/−^* mice with AIA could be TF-dependent and driven by SAA, itself induced by IL-6. In contrast to SAA, CRP levels were more variable overall in AIA than healthy mice with only a small trend towards higher levels in *IL27ra^−/−^* and *IL6ra^−/−^* strains (Supplementary Figure 3 C).

### Circulating pro-coagulant enzymatically-oxidized phospholipids (eoxPL) are increased in murine AIA

Cagulation initiated by TF requires a pro-coagulant membrane provided by circulating cells and extracellular vesicles. EoxPL are a central component of this, generated by blood cells following activation ^29,30^, but whether they are elevated in AIA, and contribute to thrombotic risk is unknown. A panel of eoxPL including isomers generated by LOX or COX enzymes in platelets and white blood cells (WBCs) were measured in mouse blood cells using lipidomics. These include the following: 12-HETE-PEs (platelets, *Alox12*; leukocytes, *Alox15*), 11-HETE-PEs (platelets or leukocytes, *Ptgs1*), 15-HETE-PEs (leukocytes, *Alox15*; platelets or leukocytes, *Ptgs1*), 5-HETE-PEs (leukocytes, *Alox5*) and 8-HETE-PEs (non-enzymatic or a minor product from *Alox12*)^31^. First, it was confirmed that immunization alone didn’t significantly change blood cell eoxPL levels (Supplementary Figure 4). The most abundant were 12-HETE-PEs, including both plasmalogen and diacyl species with fatty acyls (FA) 18:0, 18:1 or 16:0 at *Sn1* (Supplementary Figure 5 A). 12-HETE-PEs generated by platelet 12-LOX (*Alox12*) or leukocyte 12/15-LOX (*Alox15*) were the most quantitatively abundant positional isomers (Figure 1 D-F, Supplementary Figure 5 A-C).

### The AIA-induced increase in HETE-PL is dependent on enzymatic oxidation and inflammation/IL-6

EoxPL generation in AIA was compared for WT and genetically-altered strains. Similar to SAA and TATs, total HETE-PEs were elevated in blood cells from WT and *Il27ra^−/−^* mice, most significantly at day 10, while they were not increased in *Il6ra^−/−^* mice at either timepoint (Figure 1 C). This was mainly driven by 12-HETE-PEs, with the same pattern seen for all individual groups of positional isomers, although the difference was not significant for 5-HETE-PEs (Figure 1 D-F, Supplementary Figure 5 B,C). Next, to determine whether HETE-PEs were enzymatically generated, lipid extracts were saponified, then free HETEs analyzed for *S*- and *R*-enantiomer composition. More than 85% of 12-HETE was *S*-configuration, confirming generation by either 12- or 12/15-LOX (Figure 1 G). For 15-HETE, *S/R* ratio varied from 53-77 %, depending on strain and disease stage (Figure H). Notably, this isomer can be generated as the S-enantiomer by LOX, or as *S*- or *R*-enantiomers by COX, so may originate from either enzyme in this complex tissue^32,33^. 11-HETE was < 20 % *S*-enantiomer, consistent with generation by COXs (Figure 1 I). Due to the low abundance of 8- and 5-HETEs, determining chirality of these lipids wasn’t possible. Overall, this confirms that most abundant HETE-PEs generated during AIA are enzymatically generated.

### Blood cell 12-HETE-PEs are generated by platelet 12-LOX (Alox12) in AIA, not leukocyte 12/15LOX (Alox15)

In mice, the elevated 12-HETE-PEs generated during AIA development in blood cells could originate from either 12/15-LOX *(Alox15*) or platelet 12-LOX (*Alox12*). To determine which isoform and immune cells are responsible, AIA was induced in *Alox15^−/−^* or *Alox12^−/−^*mice. Prior to AIA induction, *Alox12^−/−^* blood cells contained similar levels of most HETE-PEs to WT, apart from 5-HETE-PE which was significantly elevated (Supplementary Figure 6 A,B). This suggests that neutrophil 5-LOX may be more active in blood cells following *Alox12* deletion. In contrast, *Alox15^−/−^* blood cells contained higher levels of several HETE-PEs, including 12-, 15-, 11- and 8-HETE-PE (Figure 2 A-C; Supplementary Figure 6 A-C). This suggests higher basal platelet activity in these mice since most of these are generated by platelet 12-LOX and COX-1.

**Figure 2.**
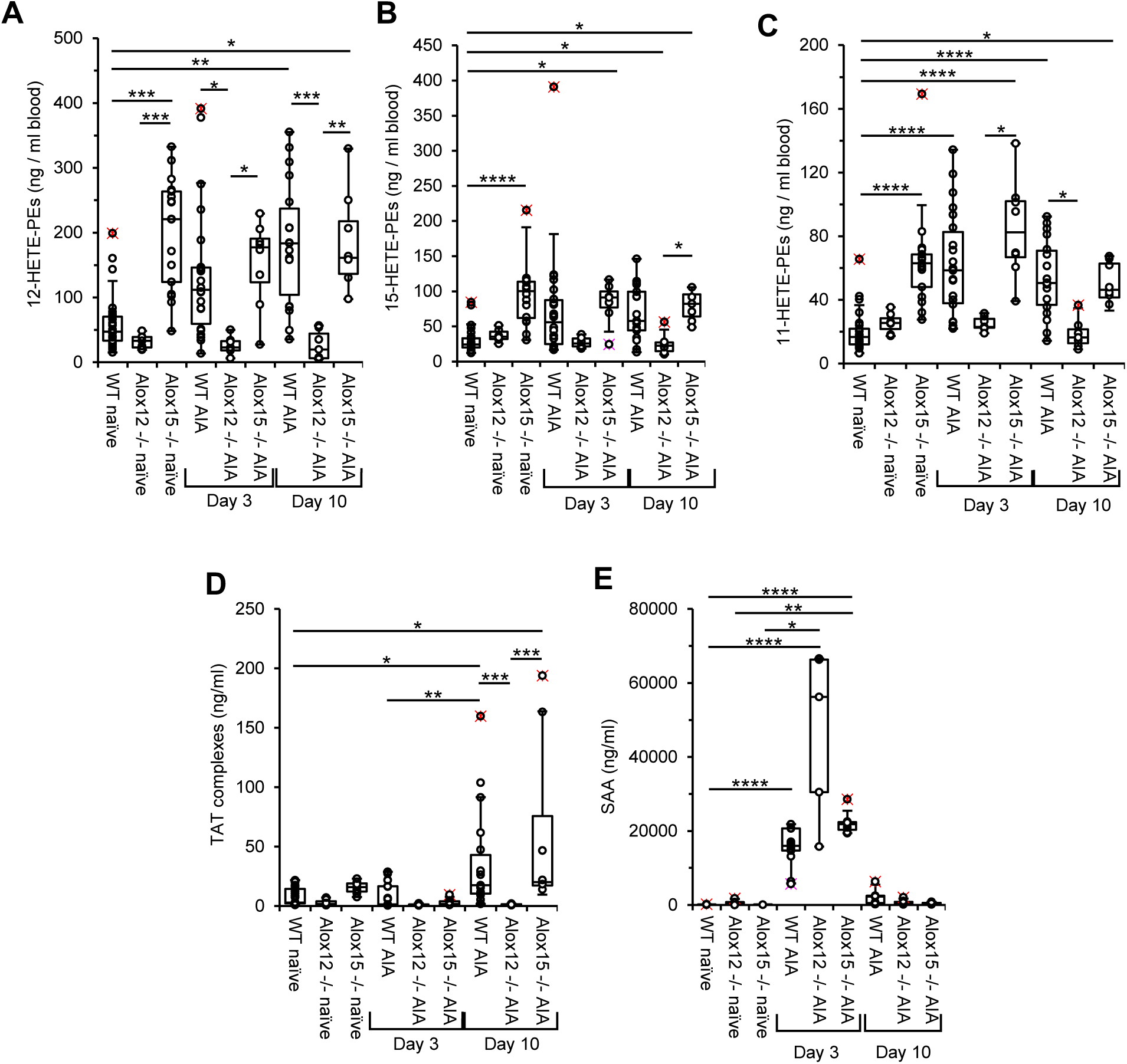
*Alox12* deletion prevents elevation of TAT complexes and generation of 12-HETE-PEs levels in mouse blood cells during AIA development, unlike *Alox15* deletion. AIA was induced in 8-12 week old *WT, Alox12*^−/−^, *Alox15*^−/−^, *Il27ra*^−/−^ and *Il6ra*^−/−^ male mice as described in Methods, with whole blood collected on Days 3 and 10. *Panel A-C. 12-, 15-, and 11-HETE-PEs were significantly elevated at day 10 of AIA, but not in Alox12^−/−^ mice.* HETE-PL were quantified as outlined in Methods using LC/MS/MS. Whole blood was collected at day 0 from WT naïve (n = 26), *Alox12^−/−^* naïve (n = 7) and *Alox15^−/−^* naïve (n = 8), as well as on days 3 and 10 of AIA development *WT* (n = 22 and 18 respectively), *Alox15^−/−^* (n = 8 for both days) and *Alox12^−/−^* (n = 7 for both days) male mice*. Panel D. TAT elevation in AIA is dependent on Alox12, but not Alox15.* TAT complexes were measured using ELISA. Plasma was collected on day 0 from WT naïve (n = 30), *Alox15^−/−^*naïve (n = 8) and *Alox12^−/−^* naïve (n = 7), as well as on days 3 and 10 of AIA development in *WT (n = 19 and 18, respectively), Alox15^−/−^* (n = 8 for both days) and *Alox12^−/−^* (n = 7 for both days) mice. *Panel E. SAA elevation in AIA is not dependent on Alox12 or Alox15.* SAA was measured using ELISA. Plasma was collected on day 0 from WT naïve (n = 9), *Alox15^−/−^* naïve (n = 4) and *Alox12^−/−^* naïve (n = 6), as well as on days 3 and 10 of AIA development in *WT (n = 15 and 7, respectively), Alox15^−/−^* (n= 7 and 4, respectively) and *Alox12^−/−^* (n = 5 and 7, respectively) mice. Data is represented in box and whisker plot. Data were analyzed using Kruskal-Wallis test and Dunn’s multiple comparisons tests (*p<0.05, **p<0.01, *** p<0.001, **** p<0.0001).

Following AIA induction, the increase in most HETE-PEs seen in WT was not observed in *Alox15^−/−^*blood cells, since eoxPL levels were already increased basally in this strain (Figure 2 A-C, Supplementary Figure 6 A-C). Indeed, during AIA, most HETE-PEs were maintained in *Alox15^−/−^* blood cells at the elevated levels seen in WT mice with AIA (Figure 2 A-C, Supplementary Figure 6 A-C). This indicates that 12/15-LOX is not the source of eoxPL that become elevated in WT mice during AIA. In contrast, for *Alox12−/−* mice, there was no increase in 12-, 15-, 11- or 8-HETE-PEs, indicating that most of these either originate from, or are dependent on *Alox12* (Figure 2 A-C; Supplementary Figure 6 A-C). Considering that 15- and 11-HETE-PEs are generated by COX-1 in platelets, it is likely that the deletion of *Alox12* has a knock-on impact on secondary platelet activation. In summary, most eoxPL that are elevated in whole blood cells in AIA originate from platelets and depend on 12-LOX. Importantly, eoxPL elevation in AIA was entirely dependent on IL-6 indicating a direct link between inflammatory signaling and downstream platelet activation.

### Platelet derived oxylipins are elevated by IL-6 in AIA

Next, free oxylipins were analyzed in blood cells during development of AIA. The data showed a platelet activation signature on days 3 and 10, dominated by strong elevations in 12-HETE and 14-HDOHE, which was fully dependent on both IL-6 and platelet 12-LOX, but did not require leukocyte 12/15-LOX (See Supplementary Figures 7, 8). This further implicates IL-6 in driving platelet activation in AIA.

### Deletion of Alox12 in AIA prevents thrombin activation, but not inflammation, while deletion of Alox15 has no impact

Next, to determine whether LOX isoforms contribute to thrombosis and inflammation, plasma TATs and SAA were determined in *Alox15^−/−^* or *Alox12^−/−^* mice. *Alox12* deletion totally prevented TAT increases on day 10, while *Alox15^−/−^*mice exhibited similar levels to WT (Figure 2 D). In contrast, SAA was increased in both strains at day 3, similar to *WT* indicating that inflammation is not dependent on either LOX (Figure 2 E).

### Alox15 or Alox12 deletion worsens AIA in mice

Next, the role of *Alox15* and *Alox12* in joint pathology was determined. First, antibody titers to mBSA were not altered by deletion of either LOX, indicating that disease induction was normal (Supplementary Figure 9). In contrast, both showed more rapid development of joint swelling that resolved more slowly than for WT (Figure 3 A). Next, joints were histologically scored for arthritis index. Deletion of *Al*ox*12* resulted in a worse phenotype at day 3, while *Alox15^−/−^* mice were not different to WT (Figure 3 B, Supplementary Table 3, Supplementary Figure 10). In contrast, in *Alox15^−/−^* mice, a significant increase in subsynovial inflammation was observed at day 10, while Alox12^−/−^ were similar to WT (Figure 3 C). On day 3, synovial exudate, pannus formation and cartilage and bone erosion in *Alox12^−/−^* were at the higher end of WT values, while *Alox15^−/−^*joints were at the lower end, although none were significantly different to WT. (Figure 3 D-F). These results somewhat reflect the SAA results, where *Alox12^−/−^* mice have higher levels than both WT and *Alox15^−/−^* at day 3 of AIA (Figure 2 E). Overall, they indicate that both strains have a somewhat worsened phenotype.

**Figure 3.**
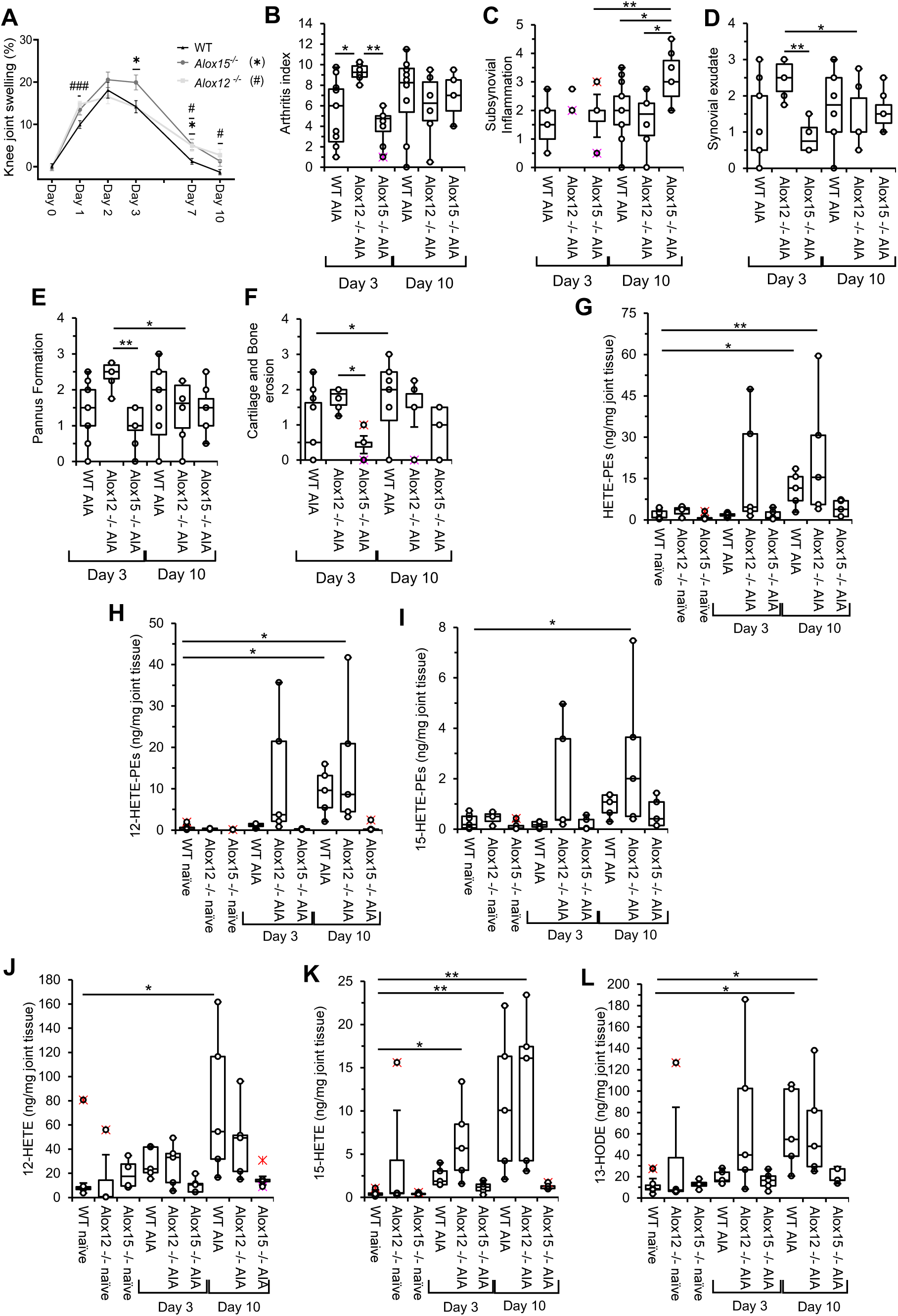
*Alox15^−/−^ and Alox12^−/−^* mice display a more severe AIA phenotype than WT, while *Alox15^−/−^* display an overall reduction of eoxPL and oxylipins in synovial tissue during AIA development. AIA was induced in 8-12 week old *WT, Alox12*^−/−^ and *Alox15*^−/−^ male mice as described in Methods, with synovial tissue collected on days 3 and 10. *Panel A. Joint swelling is increased upon Alox12 and Alox15 deletion.* Knee diameter was measured every 1-2 days and knee joint swelling was calculated as a percentage relative to knee diameter at day 0 (before arthritis induction) in WT (n = 44), *Alox15^−/−^*(n = 32) and *Alox12^−/−^* mice (n= 14). Data represent mean ± SEM, and statistical analysis was performed using a two-way ANOVA and Mix-effect analysis, comparing, on different days of AIA development, WT with *Alox15^−/−^* (*p<0.05) and WT with *Alox12^−/−^* mice (#p<0.05, ###p<0.001). *Panels B-F. Arthritis index in AIA is more severe upon Alox12 deletion.* Knee joints were collected on day 3 and 10 from WT (n = 11 for day 3; n = 12 for day 10), *Alox12^−/−^* (n = 6) and *Alox15^−/−^* mice (n = 8 for day 3 and n = 7 for day 10) of AIA for histological staining and assessment, as described in Methods. Histopathology scoring of AIA was used to obtain Arthritis index, subsynovial inflammation, synovial exudate, pannus formation, and cartilage and bone erosion. Data were analyzed using two-way ANOVA test and Šídák’s multiple comparisons test, according to time point (*p < 0.05, **p < 0.01). *Panels G-I. HETE-PE generation in the synovial tissue during AIA is dependent on Alox15.* Total HETE-PEs, 12- and 15-HETE-PEs, analyzed using LC/MS/MS, as outlined in Methods are shown. Joints were collected and pooled at day 0 from WT naïve (n = 6), *Alox12^−/−^* naïve (n = 4) and *Alox15^−/−^* naïve (n = 5), as well as on days 3 and 10 of AIA development *(n = 5 for both days). Panel J-L. Generation of 12- and 15-HETE during AIA is dependent on Alox15.* 12- and 15-HETE, along with 13-HODE, were quantified using LC/MS/MS, as outlined in Methods. Joints were collected and pooled at day 0 for WT naïve (n = 6), *Alox12^−/−^* naïve (n = 4) and *Alox15^−/−^* naïve (n = 5), as well as on days 3 and 10 of AIA development (n = 5 for each). Data are represented in box and whisker plots, displaying individual values. Data were analyzed using Kruskal-Wallis test and Dunn’s multiple comparisons test (* p <0.05, ** p <0.01, *** p<0.001, **** p<0.0001).

### Alox15 deletion prevents 12-HETE-PE generation in synovial tissue during AIA, while deletion of Alox12 has no impact

Next, the profile of eoxPL in knee joints was analyzed to determine their temporal generation and biochemical origin. Basal eoxPL levels in synovial tissue were overall quite low and not different for *Alox15^−/−^*or *Alox12^−/−^* mice, compared to WT (Figure 3 G, Supplementary Figure 11 A). On induction of AIA in WT mice, HETE-PEs significantly increased in joints on day 10 (Figure 3 G, Supplementary Figure 11 A). This was mainly due to increased levels of 12-HETE-PEs, which were the most abundant isomers detected (Figure 3 H, Supplementary Figure 11 A). In comparison, other HETE-PEs were detected at low levels (Figure 3 I, Supplementary Figure 11 A,B). Strikingly, no increase in 12-HETE-PEs was seen for *Alox15^−/−^* joints upon AIA development, confirming their enzymatic origin. Synovial tissue from *Alox12^−/−^*mice with AIA showed a significant but highly variable increase in 12-HETE-PEs at day 10 (Figure 3 H). 15-HETE-PE behaved similarly to 12-HETE-PE (Figure 3 I). These lipids can be generated by tissue macrophages expressing 12/15-LOX in mice, suggesting its expression/activity may be increased in AIA, when *Alox12* is deleted. Other HETE-PEs were detected at very low levels and were not significantly impacted by AIA (Supplementary Figure 11 A,B). Overall, in knee joints, the dominant source of 12-HETE-PEs generated in response to AIA induction is *Alox15*, contrasting with the primary source of these lipids in blood during AIA, which was *Alox12*.

### Alox15 deletion reduced generation of multiple oxylipins in synovial tissue during AIA development

Free oxylipins were next analyzed in synovial tissue from WT, *Alox15^−/−^* and *Alox12^−/−^* mice following arthritis induction. No significant differences were observed at baseline (Supplementary Figure 12). During AIA, WT joints showed substantial elevations in 12-, 15-HETE and 13-HODE, indicating 12/15-LOX activity (Figure 3 J-l; Supplementary Figure 12 A). 12- and 15-HETE were not elevated in *Alox15^−/−^* joints (Figure 3 J,K). 5-HETE was significantly elevated in WT mice on day 3, and this was not observed in either LOX-deficient strain, however, levels were very low overall (Supplementary Figure 12 C). PGE_2_ was significantly increased in *Alox15^−/−^* joints at day 10 of AIA, while PGD_2_ was not detected in synovial tissue from *Alox12^−/−^* mice (Supplementary Figure 12 E,F). Some other PGs and LTs were detected at very low concentrations, even after AIA induction (Supplementary Figure 12 A). 7,17-diHDOHE was detected in WT joints at day 10, however, the level was below the assay LOQ with insufficient quantities to allow chiral analysis for ResolvinD5 structure verification^34^. No other specialized pro-resolving candidates were detected at any timepoint. Overall, these data show that *Alox15* is the primary source of joint-derived 12- and 15-HETE in AIA-driven synovitis.

### RA patients have higher TATs and platelet counts, but similar white cell counts to healthy controls (HC)

Our murine studies demonstrated that AIA is associated with increased thrombosis, evidenced by TAT elevations driven by platelet 12-LOX in response to inflammation/IL-6. Next, we determined whether the thrombotic tendency in human RA is similarly linked with platelet 12-LOX. For this, blood was obtained from patients and HC (Supplementary Table 1). TAT complexes were significantly elevated in RA patient plasma, demonstrating increased thrombin generation *in vivo* (Figure 4 A). Also, RA patients displayed a significantly higher platelet count than HC, but no change in WBC numbers (Figure 4 B,C). These data confirm this cohort is experiencing elevated coagulation.

**Figure 4.**
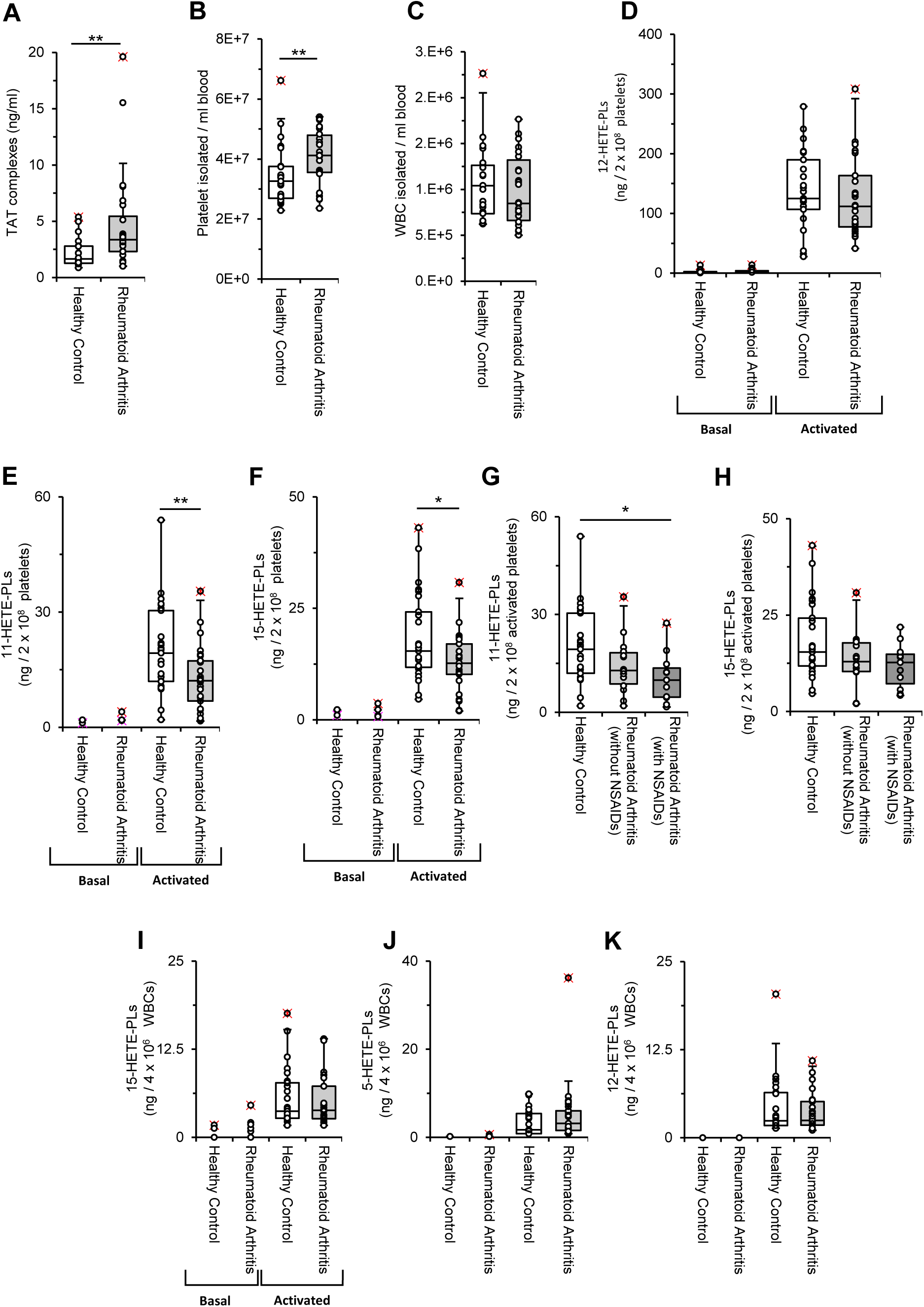
TATs and platelets are elevated in RA, while eoxPL from platelets are somewhat reduced in RA due to NSAID supplementation. *Panel A. TAT complexes are increased in RA patients compared to HC.* TAT complexes were measured using ELISA. Plasma was collected as described in Methods from RA (n = 24) and HC (n = 19). Data was analyzed using Mann-Whitney test (**p <0.001). *Panel B. RA patients have increased platelets in circulation than HC.* Platelets were isolated from HC (n = 23) and RA (n = 26) as described in Methods, then counted and calculated to provide the equivalent of 1 ml blood. Data was analyzed using Mann-Whitney test (**p <0.001). *Panel C. WBC numbers are similar between RA patients and HC.* WBC were isolated, as described in Methods, from HC (n = 25) and RA (n = 26). Cells were counted and calculated to provide the equivalent of 1 ml of blood. Data was analyzed using Mann-Whitney test. *Panel D-F. 11- and 15-HETE-PL, but not 12-HETE-PL, are decreased in RA platelets upon activation compared to HC.* Washed platelets from RA (n=25) and HC (n=23) were analyzed for eoxPL by LC/MS/MS. Platelets were activated with 0.2 U/ml thrombin and 1 mM CaCl_2_, as described in Methods. Washed platelets, either basal or activated, were analyzed for eoxPL by LC/MS/MS. Data were analyzed using Students t-test between activated conditions (*p<0.05,**p<0.01, ***p<0.001). *Panel G-H. 11-HETE-PL is decreased in activated platelets in RA due to NSAIDs.* Washed platelets from RA taking NSAIDs (n = 11), or not (n = 14), along with HC (n = 25) were isolated and activated with 0.2 U/ml thrombin and 1 mM CaCl_2_, as described in Methods. 11- and 15-HETE-PL were analyzed using LC/MS/MS. Data were analyzed using one-way ANOVA and Tukey’s multiple comparison test (*p<0.05,**p<0.01). *Panel I-K. 15-, 5- and 12-HETE-PL in WBCs are similar between RA and HC.* WBCs were isolated from RA (n = 25) and HC blood (n = 25). OxPL in resting and ionophore activated WBCs were analyzed by LC/MS/MS as described in Methods. Data were analyzed using Students t-test between activated conditions.

### RA patients have similar 12-HETE-PL profiles to HC but reduced 11- and 15-HETE-PL due to NSAID consumption

Next, washed platelets were isolated, and analyzed for eoxPL, either basally or following thrombin activation. Basal levels may better reflect circulating amounts, while activation demonstrates their capacity for generation of eoxPL. Here, HETE-PCs were also analyzed since they have been detected in human platelets^30^. Washed platelets were resuspended at a typical circulating blood count (2 × 10^8^ cells/ml), prior to stimulation and lipid extraction. Low/absent levels of HETE-PL were found basally, with only PC 16:0a_12-HETE and PC 18:0a_12-HETE species detected in more than 50 % of the analyzed patient samples. The sum of these 12-HETE-PC species displayed similar levels for RA and HC platelets, when expressed on a per cell basis (Supplementary Figure 13 A).

As expected, thrombin significantly increased 12-, 11- and 15-HETE-PL in platelets (Figure 4 D-F). While RA patient’s platelets generated similar levels of 12-HETE-PL, lower levels of 15- and 11-HETE-PL were detected than in HC platelets (Figure 4 E,F). Since these isoforms are generated by COX-1, and 30 % of RA patients were on NSAIDs, we next tested the impact of these medications (Supplementary Table 1). A small non-significant reduction was seen in RA platelets from patients not using NSAIDs, which this was further suppressed by NSAIDs, significantly for 11-HETE-PLs (Figure 4 G,H). Overall, considering that 12-HETE-PL were the most abundant generated, and their generation wasn’t impacted by aspirin (Supplementary Figure 13 B), this small reduction in COX-derived isomers did not lead to significant reductions in total eoxPL in RA (Supplementary Figure 13 C). Since RA was associated with thrombocytosis, we predict that an overall higher level of eoxPL in the circulation maybe experienced by patients if platelets become activated in disease and this will be tested for by measuring immunoreactivity to HETE-PEs.

### WBC from RA patients generate similar eoxPL levels to HC

Next, eoxPL generation by WBCs was determined for RA and HC. Basally, only PC 16:0a_8-HETE was detected in >50 % of samples, with all other HETE-PLs undetected (Supplementary Figure 14 A). Following ionophore activation, several HETE-PL were generated, while PC 16:0a_8-HETE was not detected (Supplementary Figure 14 B). No significant differences were observed between RA and HC groups for any positional isomers (Figure 4 I-K). Formation of 15- and 5-HETE-PL (Figure 4 I,J) is consistent with 5-LOX and 15-LOX in neutrophils and eosinophils, respectively. The presence of 12-HETE-PL suggests some possible platelet contamination (Figure 4 K). Overall, these data indicate that WBC eoxPL are unlikely to contribute to the pro-thrombotic tendency in RA.

### Plasma from RA patients have elevated anti-HETE-PL IgG immunoreactivity

Levels of eoxPL in platelets significantly increased during early AIA in mice. Considering that RA patients have a chronic illness, and are on several immunosuppressive medications, eoxPL generation during early disease could have been missed or may be suppressed due to treatment. To test whether RA patients might have been chronically exposed to eoxPL as part of the disease, we determined their serum IgG response to these lipids. Here, immunoreactivity to HETE-PEs was examined in serum samples from RA patients and HC (Supplementary Table 2). Immunoreactivity towards the unoxidized form, PE 18:0a_20:4 (SAPE) was first compared with that directed against 5 positional isomers of PE 18:0a_HETE-PE, including both platelet and leukocyte isoforms. First, it was noted for HC serum, that there was a higher immunoreactivity against almost all HETE-PE isoforms, than for SAPE (Figure 5). For 12-,15- and 11-HETE-PE, this was significantly elevated (Figure 5 A-C). This suggests that even in healthy subjects, exposure to eoxPL drives an immune response evidenced by elevated reactive circulating IgG. The IgG response to SAPE was not different between RA and HC serum, however, RA serum was more immunoreactive towards all HETE-PEs analyzed than HC, with 12-, 15- and 8-HETE-PEs being significantly higher in patients (Figure 5 A,B,D). Overall, these data indicate that RA patients have a higher antigenic response to HETE-PEs than healthy controls, suggesting higher chronic exposure to these lipids *in vivo*, in particular 12- and 15-HETE-PEs, both of which are generated by platelets.

**Figure 5.**
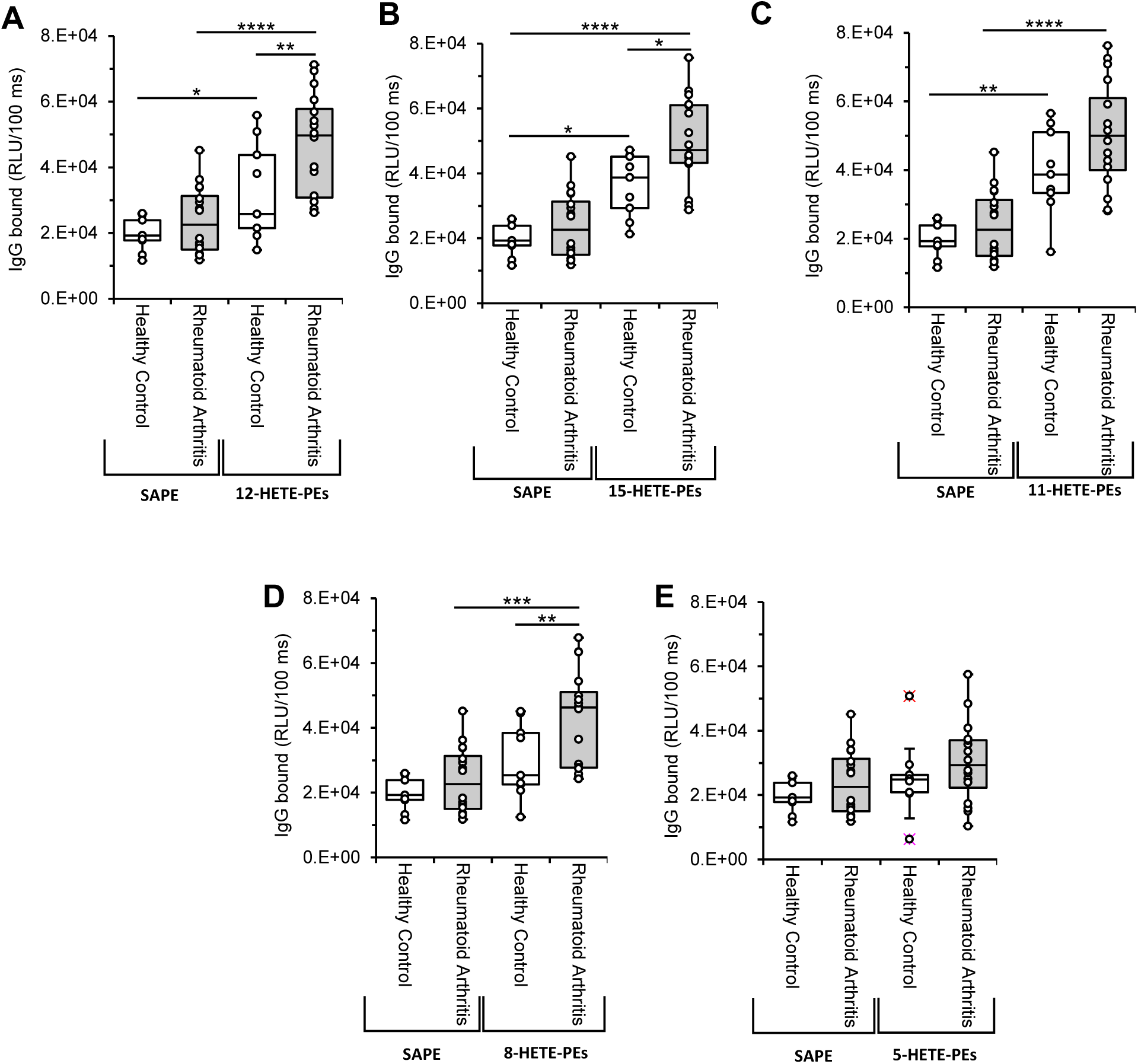
Circulating IgG recognizing HETE-PEs are increased in RA. *Panels A-E. IgG directed against 12-,15- and 8-HETE-PEs are significantly increased in RA patients compared to healthy controls.* IgG levels against 12-, 15-, 11-, 8- and 5-HETE-PEs were determined as outlined in Methods and compared to IgG levels against SAPE (the same levels in all figures) in serum from HC (n=9) and RA (n=16) as described in Methods. Data were analysed using two-way ANOVA and Šídák’s multiple comparisons tests (*p<0.05, **p<0.01, ***p<0.001, **** p<0.0001).

## Discussion

RA patients are at higher risk of thrombosis, often leading to mortality^3,4,35^. Here, we reveal using genetically-altered mice and human cohorts that platelet *Alox12* drives this, at least in part through bioactivity of its pro-coagulant eoxPL enhancing thrombin generation^8,12,36^. A critical role for IL-6 in driving this phenomenon in mice was found.

Herein, we applied an established AIA model to WT^25,37^, *Il27ra^−/−^* ^38^ and *Il6ra^−/−^* ^39^ mice, where each develop specific synovial pathotypes of RA, with features similar to human disease^26,40^. The increase in HETE-PEs seen in WT and *Il27ra^−/−^* mice with AIA was mainly driven by 12-HETE-PEs, suggesting platelet involvement, and this was subsequently confirmed using *Alox12^−/−^* mice. The cellular origin of AIA-driven eoxPL in mouse blood cells was exclusively platelet 12-LOX, with no contribution of the leukocyte isoform. This agrees with a previous study showing that generation of eoxPL in *ex vivo* clotting whole blood from healthy WT mice is mainly *Alox12*-dependent^21^. Furthermore, mice lacking *Alox12* did not show elevated TATs in AIA, directly linking coagulation activation *in vivo* with this isoform. Platelet involvement was further implicated by elevations in 15- and 11-HETE-PEs, generated by platelet COX-1.

Both WT and *Il27ra^−/−^* mice developed leukocyte-driven synovitis and systemic inflammation, as evidenced by increased circulating SAA ^23^. Alongside this, they showed significantly increased TAT complexes at day 10. On the other hand, *Il6ra^−/−^* mice did not demonstrate increases in blood cell eoxPL, SAA or TAT complexes. These mirror clinical findings in human patients treated with tocilizumab, an antibody against IL6-R, that controls inflammation by inhibiting the expression of acute phase reactants, including fibrinogen^41^. Furthermore, IL-6 was previously shown to increase TATs in human plasma ^42^. Taken with our findings, this reveals that IL-6 likely directly drives eoxPL generation and downstream coagulation in AIA.

In contrast to our finding that platelet 12-LOX is exclusively responsible for blood cell eoxPL and TAT elevations, the leukocyte isoform only generated eoxPL in joints during AIA, regulating influx of inflammatory cells into the synovium. Indeed, eoxPL can induce an anti-oxidative response^43^, and oxylipins such as 12-HETE, 15-HETE and 13-HODE, which were significantly reduced in synovial tissue on *Alox15* deletion, are also known PPARγ ligands^44^. This suggests that 12/15-LOX is protective in synovitis, consistent with results from other mouse models of arthritis, through mechanisms that could involve both eoxPL and free oxylipins^45^. Importantly, the data show that the two LOX isoforms play distinct tissue-specific roles in AIA responsible for different pathological outcomes. Neither LOX-deficient strain showed any reduction in SAA, indicating that inflammation acts upstream of eoxPL/TAT generation in AIA.

Consistent with previous reports, patients with RA displayed thrombocytosis and increased TATs^14,46^Although higher circulating eoxPL weren’t observed (as cell counts were normalized prior to measuring the lipids), the elevated immunogenic response suggested chronic exposure in the circulation of patients. This could be due to an increase in platelet counts in circulation, which we also observed in our cohort, or via higher eoxPL generation during disease development. In agreement with our findings, elevated plasma IgG directed against eoxPL was previously shown in another pro-thrombotic disorder, anti-phospholipid syndrome^12^.

Despite increased thrombosis risk in RA, guidelines don’t currently recommend long-term prophylactic anticoagulation, with patients following the same rules as the general population^47,48^. In murine AIA, increased eoxPL, SAA and TATs were completely dependent on IL-6, while *in vitro* SAA induces TF expression and activity^49^. This suggests that biological medicines or small molecule inhibitors that target IL-6 signaling may reduce thrombosis in patients both through reducing TF as well as eoxPL. As evidence, a cohort study found reduced coagulation biomarkers in early RA following antirheumatic treatment with anti-cytokine targeting therapies (e.g. tocilizumab, certolizumab) as compared to conventional DMARDs (e.g. methotrexate)^41^.

In conclusion, LOX isoforms play distinct cell and tissue-specific roles in driving either local or systemic pathology of AIA. Platelet 12-LOX is a central contributor to the higher thrombotic risk in AIA, through supporting the generation of pro-coagulant membranes. In human RA, higher immunoreactivity to HETE-PEs combined with higher platelet counts (and robust generation of eoxPL upon thrombin challenge) suggest that similar to mice, platelet 12-LOX also directly contributes to thrombotic risk. These data indicate that reducing inflammation using DMARDs, particularly those targeting IL-6, should also suppress platelet reactivity and improve thrombosis risk in RA.

## Authorship Contributions

DOC, STOH, RHJ, ACF, GWJ, JJB, BM, FM, DH, ASM and CG performed the research. RHJ, ACF and GWJ provided training for the AIA mouse model. DOC, VJT, MBP, AAH and VBOD contributed to the methodology. EC supported the Human clinical cohort (Cardiff) study, which provided essential samples from RA patients. AB and REMT supported the immunological clinical cohort (Leiden), which provided serum samples of RA patients. MG, PVJ, PWC, SAJ, VBOD provided conceptualization. SAJ and VBOD provided supervision. DOC, SAJ and VBOD drafted and edited the manuscript. All authors edited the manuscript.

## Data Sharing Statement

All processed data is included in a supplemental file, while raw data is available on reasonable request.

## Novelty and Significance

### What is known?

1. Thrombosis is a major cause of morbidity and mortality in RA.
2. The mechanisms underlying the increased thrombotic risk in RA are unknown.
3. Patients with RA are not currently treated to reduce thrombotic risk.

### What new information does this article contribute?

This study investigated the role of the membrane surface of circulating platelets and white cells in driving the thrombotic risk commonly associated with RA in humans and animal models, and how this might be modulated by inflammation caused by the disease. We found that induction of arthritis in mice led to increased circulating coagulation markers along with generation of pro-coagulant lipids in platelets. Using genetically-modified mice, we demonstrated the lipids were generated by a platelet enzyme, 12-lipoxygenase (12-LOX, Alox12) in an interleukin-6 (IL-6) dependent manner. Thrombotic risk was completely prevented if either 12-LOX or IL-6 signaling were inhibited. In humans, we found that RA patients showed higher levels of thrombosis markers, as well as evidence of long-term exposure to the same pro-coagulant lipids, as measured by immunoreactivity assays. Our study indicates that targeting IL-6 using DMARDs, which are already prescribed in RA to dampen inflammation, may also reduce the risk of thrombotic complications. This is the first characterization of the procoagulant surface of platelets in arthritis, with the membrane itself representing a potential new target for anti-coagulant therapies.

## Supplementary Methods, Results and Data

### Supplementary Methods

#### Human clinical cohort (Cardiff)

Human experiments were performed under the project titled “Cardiff Regional Experimental Arthritis Treatment and Evaluation Centre” approved by the Ethics Committee for Wales (Reference No 12/WA/0045) and performed in accordance with the principles of the Declaration of Helsinki, with informed consent and with full ethical approval. Rheumatoid arthritis patients were recruited for venous blood sampling during a routine appointment. Patients had no history of venous and/or arterial thrombosis at time of venipuncture. Sex-matched healthy volunteers were recruited, excluding those with history of high cholesterol, arterial or venous thrombosis, recurrent fetal loss, cardiac disease, diabetes, or any chronic inflammatory disease. The average age of healthy volunteers was 51 years-old. Healthy control individuals were instructed to not take aspirin, non-steroidal anti-inflammatory drugs, or other medications for 14 days before blood donation. Recruitment took place between February 2020 to April 2022, recruiting a total of 25 gender-matched healthy volunteers, and 26 RA patients. Clinical characteristics of recruited subjects are in Supplementary Table 1. Sample size was determined based on data from a previous study ^1^, where using an online sample size calculator^2^, each study group needed at least 25 participants to achieve statistical significance. Outliers were included. For TAT complexes, 6 HC and 2 RA plasma samples were not determined for lack of enough volume to complete the analysis. For platelet count experiment, 2 healthy control platelet samples partially clotted, preventing an accurate comparable count. For platelet and WBC eoxPL analysis, 1 RA sample was not analyzed due to low blood volume obtained during collection.

**Supplementary Table 1.** Baseline clinical characteristics of recruited volunteers in the clinical cohort. (CCP: Cyclic Citrullinated Peptide, CRP: c-reactive protein, DAS: disease score activity, p-value tests: Fisher exact for categorical (a) or Mann-Whitney test for continuous variables (b).

| **Variable** | **Healthy control**<br>**(HC) [n=25]** | **Rheumatoid Arthritis**<br>**(RA) [n=26]** | **p** |
| --- | --- | --- | --- |
| **Age (mean ± SD)** | 51.69 ± 8.74 | 61.08 ± 16.51 | 0.0224 (b) |
| **Female sex** | 21 (84%) | 23 (88%) | 0.6514 (a) |
| **DAS28 (mean ± SD)** | – | 2.68 ± 1.45 | – |
| **Disease duration (years)**<br>**(mean ± SD)** | – | 7.58 ± 8.43 | – |
| **Rheumatoid Factor (+)** | – | 60% | – |
| **Anti-CCP (+)** | – | 64% | – |
| **Erythrocyte sedimentation rate**<br>**(mm/hour ± SD)** | - | 14.12 ± 12.9 | - |
| **CRP (mg/L, mean ± SD)** | - | 4.8 ± 7.37 | - |
| **Aspirin use** | 0 | 8% | – |
| **NSAIDs use** | 0 | 30% | – |
| **Smoker** | 0 |  | – |
| **Osteoarthritis** | 0 | 23% | – |
| **Hypertension** | 0 | 19% | – |
| **Diabetes** | 0 | 0 | – |
| **Hypothyroidism** | 0 | 12% | – |
| **Statin use** | 0 | 0 | – |

#### Washed platelet isolation from human blood

Human blood was collected using a 21-gauge butterfly needle and two 20 ml venipuncture syringes. Each syringe contained 3.6 ml ACD [2.5% (w/v) trisodium citrate, 1.5% (w/v) citric acid, and 100 mM Glucose], and a final volume of 18 ml was obtained. The blood was centrifuged without brake (400 g, 10 mins, 22°C). The platelet-rich plasma (PRP) was isolated and recentrifuged without brake (1400 g, 8 mins, 22 °C). The platelet-poor plasma (PPP) supernatant was isolated for further analysis. The platelet pellet was re-suspended using 10 ml ACD: Tyrode’s buffer [145 mM NaCl, 12 mM NaHCO_3_2.95 mM KCl, 1 mM MgCl_2_, 10 mM HEPES and 5 mM glucose] (1:9 v/v), before centrifuging (1400 g without brake, 8 mins, 22 °C). Platelets were then isolated and resuspended in 1 ml Tyrode’s buffer. Platelets were counted with a hemocytometer and resuspended at 2 × 10^8^ cells per ml in Tyrode’s buffer. A total of 3 × 10^8^ platelets were used either as unstimulated controls, or stimulated with 0.2 U/ml thrombin and 1 mM CaCl_2_ for 30 minutes at 37 °C.

#### White blood cell isolation from human blood

Human blood was collected using a 21-gauge butterfly needle and a 60 ml syringe. 20 ml blood was drawn into 4 ml Hetasep™ (Stem Cell Technologies, France) and 4 ml 2 % Citrate. After gentle inverting to mix, the syringe was left in an upright position for 1h to achieve separation. The top layer was isolated and centrifuged (400 g without brake, 10 mins, 4 °C). The pellet was resuspended in DPBS/0.4 % citrate, followed by centrifugation (400 g, 6 mins, 4 °C). 5 ml 0.2 % hypotonic NaCl was added to lyse red blood cells (RBCs), followed by a washing step with 50 ml ice-cold DPBS/0.4 % Citrate. The WBCs were centrifuged (400 g, 6 mins, 4°C) and a second RBC lysis was performed. After pelleting, WBC were resuspended in 1 ml Krebs buffer [0.1 M NaCl, 5 mM KCl, 47.7 mM HEPES, 1 mM NaH_2_PO_4_·2H_2_O, 2 mM glucose, pH 7.4] and counted using a hemocytometer. Cells were then resuspended at 4 × 10^6^/ml. A total of 6 × 10^6^ WBCs were used as unstimulated controls, or were stimulated using 10 μM A23187 and 1 mM CaCl_2_ for 30 minutes at 37 °C. Lipids were then extracted as described below.

#### EoxPL extraction

To each sample, 2.5 ml hexane: IPA:1 M acetic acid (30:20:2 v/v), was added, with 10 ng DMPE [PE 14:0/14:0] and DMPC [PC 14:0/14:0] as internal standards. After 30 mins incubation on ice, 2.5 ml hexane was added, followed by vortexing and centrifuging (1475g, 5 mins, 4 °C). The top layer was recovered and placed at 4 °C. 2.5 ml of hexane was added to the bottom layer, followed by vortexing and centrifuging again (1475g, 5 mins, 4 °C). The top layer was recovered and combined with the previously isolated top phase, then dried using a Rapidvap N2/48 (Labconco Corporation). Lipids were resuspended in 200 µl methanol, and stored at − 80 °C under N_2_ prior to analysis by LC/MS/MS.

#### HETE-PL standards

HETE-PLs were generated as mixed isomers, and used either as a racemic mixture or as individual positional isomers once isolated and purified, as previously described^3^. Briefly, 5 mg 1-stearoyl-2-arachidonoyl-sn-glycero-3-phosphoserine (SAPC) or 1-stearoyl-2-arachidonoyl-sn-glycero-3-phosphoethanolamine (SAPE) was resuspended in 1.5 ml methanol and 64 µl 10 mM pentamethylchromanol. The mixture was dried using N_2_, then air oxidized by incubation at 37 °C for 24 hours. The mixture was resuspended in 200 μl methanol, and 10 µl 100 mM SnCl_2_ added for 10 min to reduce the lipids to PC 18:0a/HETE or PE 18:0a/HETE. Lipids were extracted by adding 3.3 ml extraction solvent [MeOH:CHCl_3_:H_2_O (8:20:5, v/v/v)] to the lipids, followed by vortexing and centrifugation (1475 g, 5 mins). Bottom chloroform layer was recovered and dried under N_2_, followed by resuspension in 200 µl methanol prior to the purification protocol. HETE-PLs were purified as a mixed isomer mixture using reversed-phase liquid chromatography on an HPLC instrument (1260 Infinity, Agilent Technologies). The column used was a Supelco Discovery C_18_ (25 cm x 4.6 mm x 5 μm). A gradient elution method with a flow rate of 1 ml/min was used: 50 % - 100 % mobile phase B (A: H_2_O, 5 mM NH_4_CH_3_CO_2_, B: MeOH, 5 mM NH_4_CH_3_CO_2_) for 15 minutes, then held at 100 % B for 20 minutes. The elute was monitored at 205 nm for unoxidized lipid substrate and 235 nm for oxidized lipids (HETE-PLs). Fractions were collected and stored at −80 °C prior to further analysis. For the isolation of individual positional isomers, a reversed-phase liquid chromatography HPLC method was performed using two Supelco Discovery C18 columns (25 cm x 21.2 mm x 5 μm) connected in series, with a gradient of 100 % mobile phase B (A: 100 % MeOH, B: 95 % MeOH, 5% H_2_0) for 200 minutes, followed by 100 % mobile phase A for 100 minutes with a total flow rate of 10 ml/min. Once purity was confirmed, HETE-PLs were quantified by spectrophotometry using absorbance at 235 nm, using ε_1mm,1cm_ = 28 absorbance units (au). Once quantified, HETE-PLs were stored at − 80 °C under N_2_ until use.

#### Analysis of serum IgG against eoxPL

Serum samples from rheumatoid arthritis patients were obtained from the Biobank Rheumatic diseases of the Department of Rheumatology, Leiden University Medical Center. The study was approved by the LUMC Biobank Ethical Review Committee and all patients gave written informed consent. Samples of serum from healthy controls were obtained under the project “Study of the lipidomic profile of blood clots from healthy volunteers” approved by the School of Medicine Ethics Review Committee, Cardiff University (REC/SREC reference No 16/02, study 10). Serum was obtained using a 21-gauge butterfly needle and a BD vacutainer® (Thermo-Fisher Scientific, UK). Blood was drawn to a final volume of 10 ml. After mixing, the blood was allowed to clot, over 30 min. Serum was isolated after centrifugation (2,340 g, 10 mins) and stored immediately at −80 °C. The Leiden samples were compared with serum generated from blood obtained from healthy volunteers from Cardiff, UK, since no healthy controls were available from Leiden. The blood draw and serum isolation process followed the same protocol for both sites. Clinical characteristics of both groups are given in Supplementary Table 2.

**Supplementary Table 2:**
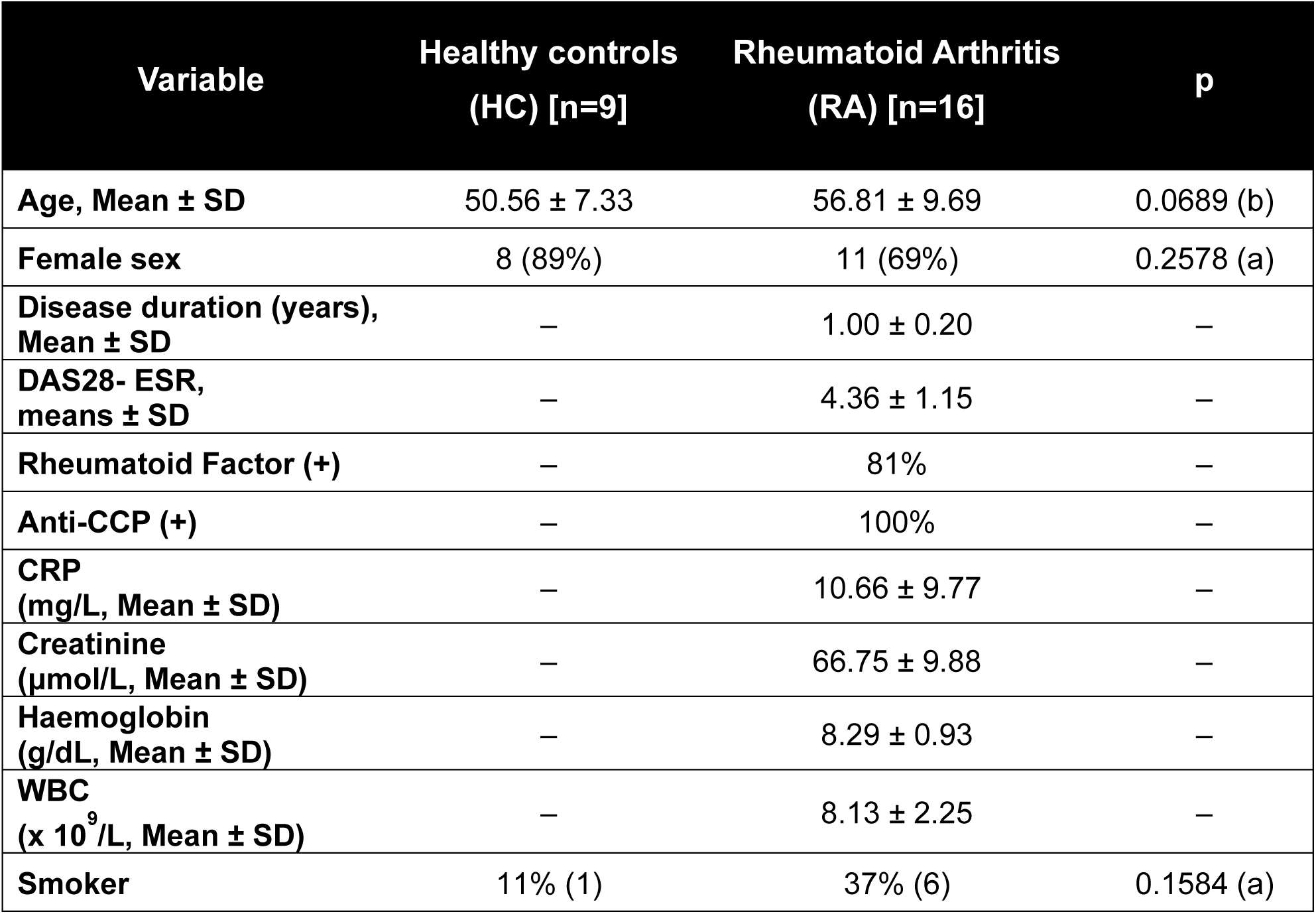
Baseline clinical characteristics of participants in the immunological clinical cohort. Data from RA patients were provided by Leiden University Medical Centre. [WCC: white cell count, ESR: Erythrocyte sedimentation rate, SD: standard deviation, p-value tests: (a) Chi-square test for categorical and (b) Mann-Whitney test.

| Variable | Healthy controls<br>(HC) [n=9] | Rheumatoid Arthritis<br>(RA) [n=16] | p |
| --- | --- | --- | --- |
| Age, Mean ± SD | 50.56 ± 7.33 | 56.81 ± 9.69 | 0.0689 (b) |
| Female sex | 8 (89%) | 11 (69%) | 0.2578 (a) |
| Disease duration (years),<br>Mean ± SD | – | 1.00 ± 0.20 | – |
| DAS28- ESR,<br>means ± SD | – | 4.36 ± 1.15 | – |
| Rheumatoid Factor (+) | – | 81% | – |
| Anti-CCP (+) | – | 100% | – |
| CRP<br>(mg/L, Mean ± SD) | – | 10.66 ± 9.77 | – |
| Creatinine<br>($\mu\text{mol/L}$, Mean ± SD) | – | 66.75 ± 9.88 | – |
| Haemoglobin<br>(g/dL, Mean ± SD) | – | 8.29 ± 0.93 | – |
| WBC<br>($\times 10^9/\text{L}$, Mean ± SD) | – | 8.13 ± 2.25 | – |
| Smoker | 11% (1) | 37% (6) | 0.1584 (a) |

#### Determination of autoantibodies against HETE-PLs

HETE-PE autoantibody titers were determined by chemiluminescent ELISA assay^1^. 5-HETE-PEs, 12-HETE-PEs, 15-HETE-PEs and 8-HETE-PEs and SAPE, were diluted to 20 µg/µl, and 25 µl added to a well of a PolySorp® surface plate (ThermoFisher scientific), followed by drying under N_2_ stream. Each well was blocked using 0.5 % (w/v) fish-gelatine in 0.27 mM EDTA/DPBS (55 µl) and incubated for 1 hour. In each well, 50 µl serum, diluted (1:12) in DPBS-0.27 mM EDTA, was incubated for 90 min. Wells were washed 3 times with DPBS/EDTA solution, before adding 25 µl anti-human IgG alkaline phosphatase-conjugated secondary antibody (Sigma Aldrich) diluted 1:20,000 in blocking solution. After another wash, 25 µl LumiPhos 530 (Lumigen, Inc), diluted 1:3 in H_2_O, was added to each well. Following incubation for 90 min, luminescence was read on a microplate reader (CLARIOstar Plus) and data expressed as relative light units/100 ms (RLU/100 ms).

#### Murine antigen-induced arthritis model

*IL27ra^−/−^* ^4^, IL*6ra^−/−^* ^5^, *IL6^−/−^*^6^, *Alox15^−/−^* ^7^ and *Alox12^−/−^* ^8^ mice were from a C57BL/6 background and were bred under specific pathogen-free conditions at Cardiff University (Cardiff, Wales). C57BL/6 wild-type (WT) mice were obtained through Charles Rivers Laboratories (UK) and went through an acclimatization period of 2-weeks prior to experiments. Mice were housed in filter top cages (or scantainers, *IL6ra−/−*), with 12-h light/dark cycles at 20– 22°C, and fed standard chow with unrestricted water. For tissue harvesting, mice were killed via CO2 inhalation, followed by cardiac puncture. All animal experiments were conducted in accordance with the United Kingdom Home Office Animals (Scientific Procedures) Act of 1986, under the project licenses PE8BCF782 and PC0174E40. AIA was induced in various strains of mice, resulting in different arthritic phenotypes similar to those observed in human RA. Induction of AIA in WT mice results in a myeloid-rich arthritis phenotype, with more diffuse immune infiltration, driven mainly by macrophages develops^9^. However, when induced in *IL27ra* deficient mice, a hyper-inflammatory phenotype, characterized by a lymphoid-rich phenotype and ectopic lymphoid-like structures develops, along with an elevated adaptive immune response^10^. In the case of *IL6ra* knockout (KO) mice, a fibroblastic-rich phenotype with reduced immune infiltration (pauci-immune) is observed^11^. Simple randomization was performed at cage level to allocate mice to the AIA model. AIA was induced in male mice, aged between 9 - 12 weeks, to minimize potential confounders, as previously described^12^. For most measures, tissue/blood from one mouse represents an experimental unit. For lipid analysis of synovial tissue, pooling of joint tissue from more than one was necessary, ranging from 3 mice (6 joints) to 6 mice (12 joints). In brief, mice were subcutaneously injected with 100 μL mBSA/Complete Freund’s Adjuvant (CFA) (Sigma-Aldrich) stable emulsion. Next, 100 μL *Bordetella Pertussis* toxin (1.6 ng/µl) was administered intraperitoneally. After one week, the mice were reimmunized with mBSA/CFA on the opposite flank. 21 days after the first immunization, inflammatory arthritis was induced using an intraarticular (i.a.) injection of 10 μL mBSA (10 mg/ml). Both i.p, s.c. and i.a. injections were performed in a simple randomized order, with the researcher blinded to genotypes. Arthritis progression was monitored by measuring knee joint swelling with a POCO 2T micrometer (Kroeplin) in the morning to reduce confounding factors. Three days after arthritis triggering, mice were culled for the acute disease time point, while the chronic time point was 10 days post i.a. injection. The sample size was initially determined based on data from a previous study^13^, using an online sample size calculator^2^ and employing TAT complexes as the primary outcome measure. Each study group was determined to require at least 8 mice. However, it was possible to reduce this number to 6 and 7 in the case of *IL6ra−/−* and *Alox12−/−* mice, respectively, since a significant difference in the primary outcome measure – TAT complexes - was observed following the first experimental group test. A total of 152 mice were used across all experiments. Animals were observed daily and monitored for signs of distress. Only males were used since females show reduced incidence and more variable phenotype in the model.

#### Mouse Blood Collection

Mouse blood collection and processing were performed as previously described ^13^. Whole blood was collected via cardiac puncture, preloaded with 100 ml of an anticoagulant mixture consisting of sodium citrate 3.8% (9:1, v/v) and 0.1 mg/ml corn trypsin inhibitor (Haematologic Technologies Inc., USA). Collected blood was centrifuged at 3000 g for 5 minutes at room temperature, followed by plasma and whole blood cells isolation. Plasma was analyzed thrombin/antithrombin (TAT), prothrombin time (PT), D-dimers, serum amyloid A (SAA).

#### TAT complexes

TAT complexes were quantified in plasma using murine TAT ELISA Kit, as per the manufacturer’s instructions (ab137994, Abcam, UK). Plasma samples were diluted at 1:100 before incubation with a TAT complex specific antibody. TAT complex specific biotinylated detection antibody was then added, followed by Streptavidin-Peroxidase conjugate. Chromogen Substrate was then added and left to react for 20 min before adding Stop Solution. Absorbance was immediately read on a microplate reader (CLARIOstar Plus, BMG Labtech) at 450 nm and values were corrected for background by subtracting readings at 570 nm. All samples and standards were analyzed in duplicate.

#### D-dimers

D-Dimers were analyzed using mouse D-Dimer, D2D ELISA Kit as per manufacturer’s instructions (CSB-E13584m, Cusabio). Plasma samples were diluted at 1:500, before adding to each well. After 2h incubation, 100 µl biotin-conjugated antibody specific for D-dimers was added. This was followed by the addition of 100 µl avidin conjugated horseradish peroxidase. Subsequently, 90 µl TMB substrate was added. After incubation with substrate, color development was stopped using a stop solution and optical density read immediately on a microplate reader (CLARIOstar Plus, BMG Labtech) at 450 nm with the background of 570 nm subtracted. All samples and standards were analyzed in duplicate.

#### Serum Amyloid A

SAA in plasma was determined using a mouse SAA ELISA Kit as per the manufacturer’s instructions (ab215090, Abcam, UK). Plasma samples were diluted 1:1000 before adding to each well. This was followed by the addition of antibody cocktail. After a 1-hour incubation, TMB Development Solution was added to each well. Stop solution was then added before reading absorbance at 450 nm on a microplate reader (CLARIOstar Plus, BMG Labtech). All samples and standards were analyzed in duplicate.

#### Prothrombin time

Prothrombin time was measured in plasma using a coagulation analyzer (Amelung KC 10). Firstly, plasma was warmed in a water bath at 37 °C for 5 min. Plasma (40 µl) was then added to plastic cuvettes with a magnetic bead and incubated at 37 °C for 5 minutes. Next, 100 µl RecombiPlasTin 2G reagent (Werfen) was added, which promotes clot formation. The forming clot entangles the magnetic bead, resulting in the rotation of the bead within the cuvette. The time between the addition of RecombiPlasTin 2G reagent and the termination of the electromagnetic coupling represents designated prothrombin time, which is measured in seconds. Samples were analyzed in duplicate.

#### Mouse whole blood cell lipid extraction

Whole blood cell pellets were resuspended in 1 ml antioxidant buffer [ice-cold DPBS, 100 μM diethylenetriaminepentaacetic acid, 100 μM butylated hydroxytoluene, 7.5 μM acetaminophen, pH 7.4], followed by the addition of 10 µl of SnCl_2_ (100 mM) for 10 min. Subsequently, internal standards were added. For whole blood lipidomics, SPLASH® LIPIDOMIX® Mass Spec Standard (Avanti Polar Lipids, USA) was used as an IS mixture, containing 10 ng PE(15:0/18:1(*d7*)) and 284 ng PC(15:0/18:1(*d7*)) added per sample. For oxylipin analysis, the IS used were deuterated lipids of the same class as the analyzed lipids, namely 2.3 ng 13(S)-HODE-d4, 2.5 ng 5(S)-HETE-d8, 2.5 ng 12(S)-HETE-d8, 2.5 ng 15(S)-HETE-d8, 2.5 ng 20-HETE-d6, 2.6 ng Leukotriene B4-d4, 2.9 ng Resolvin D1-d5, 2.5 ng Prostaglandin E2-d4, 2.7 ng Prostaglandin D2-d4, 2.7 ng Prostaglandin F2α-d4, 2.8 ng Thromboxane B2-d4, 2.8 ng 11-dehydro Thromboxane B2-d4, 2.5 ng11(12)-EET-d11 (Cayman Chemical, UK), as previously described^14^. Briefly, samples were transferred to 10 ml glass vial containing 2.5 ml ice-cold methanol. Lipids were extracted by adding 1.25 ml chloroform to each sample followed by incubation on ice for 30 minutes. Then, 1.25 ml chloroform and 1.25 ml water were added, and vortexed. The samples were then centrifuged at 400 g for 5 minutes at 4 °C to obtain a biphasic solution. Lipids were recovered from the bottom chloroform layer, and an additional 2.5 ml chloroform was then added the process repeated. The chloroform layers were then pooled and dried using a Rapidvap N2/48 evaporation system (Labconco Corporation), re-suspended in methanol, and stored at − 80 °C prior to analysis by LC/MS/MS.

#### Alkaline hydrolysis of lipid extracts for chiral HETE analysis

Whole blood cell pellet lipid extracts were dried under a stream of N_2_ and resuspended in 1.5 ml IPA. Alkaline hydrolysis was accomplished by the addition of 1.5 ml 1 M NaOH, followed by a 30 min incubation at 60 °C in a dry bath incubator. Afterwards, the extracts were acidified to pH 3.0 using 150 µl 1 M HCl before re-extraction. Briefly, to each sample 3 ml hexane was added, followed by vortexing and centrifugation (1475 g, 5 min, 4 °C). The top organic layer was recovered, and another 3 ml hexane was added to the remaining bottom layer followed by vortexing and centrifugation. The top layer was recovered and combined with the previous isolated organic layer. The IS used was 12(S)-HETE-d8, which was already present in the lipid extracts, as above. The combined layers were dried using a Rapidvap N2/48 evaporation system (Labconco Corporation). Lipids were resuspended in 150 µl methanol, and stored at − 80 °C in an N_2_ atmosphere until analysis by LC/MS/MS.

#### Mouse synovial tissue lipid extraction

Synovial tissue was dissected from the joint cavity. The tissue was weighed and pooled in order to achieve at least 5 mg of tissue. Samples were then transferred to an Eppendorf tube with 0.5 ml antioxidant buffer [phosphate buffered saline, 100 μM diethylenetriaminepentaacetic acid, 100 μM butylated hydroxytoluene, 7.5 μM acetaminophen, pH 7.4]. Synovial tissue samples were homogenized in a Bead Rupture Elite® (2 cycles at 5 m/s for 20 seconds). Tissue samples were then transferred to glass vials and the remaining tissue was washed out with a further 0.5 ml antioxidant buffer, followed by the addition of internal standards, as described for mouse whole blood cell lipid extraction. The reduction of hydroperoxides was achieved by the addition of 10 µl of SnCl_2_ (100 mM), and incubation for 10 minutes on ice. Lipids were first extracted using an isopropanol/hexane method, by adding 2.5 ml hexane/isopropanol/acetic acid (30:20:2, v/v/v) extraction solution to each sample. After vortexing, 2.5 ml hexane was added, followed by another vortexing step. The separation of phases was achieved by centrifugation (400 g, 5 mins, 4 °C). The upper layer was recovered and transferred to new extraction vials. Another 2.5 ml hexane were added, followed by another round of vortexing and centrifugation. The upper phase was recovered and combined with the previously recovered layer. The remaining bottom layer was then extracted using the Bligh and Dyer method by adding 2.5ml methanol and 1.25 ml chloroform. Samples were vortexed before adding 1.25 ml chloroform and 1.25 ml water. Samples were vortexed and centrifugated (400 g, 5 mins, 4 °C), and bottom layers recovered and dried using a Rapidvap N2/48 evaporation system (Labconco Corporation), before being resuspended in methanol. To remove possible contaminating particles such as bone or cartilage, the samples were further extracted, using a final isopropanol/hexane method, which by recovering the upper layer, keeps the sedimented contaminating particles on the bottom layer. For this, 2.5 ml hexane/isopropanol/acetic acid (30:20:2, v/v/v) extraction solution was added to the lipid extract diluted in 1 ml of water. Following vortexing, another 2.5ml hexane was added. Upper layer was recovered after centrifugation (400 g, 5 mins, 4 °C). The upper layer was combined with other hexane layers. This combined hexane layers were then dried. Lipids were re-suspended in methanol and stored at −80 °C prior to analysis by LC/MS/MS.

#### Mouse knee joint histology

Whole knee joints were recovered, and following the removal of skin, knees were placed into histology cassettes and fixed in 10 % formalin for three days. This was followed by a decalcification process, where the cassettes were incubated in decalcification buffer (10 % formic acid in water). The tissue was then processed using the HistoCore PEARL before being embedded into paraffin blocks through Arcadia H instruments (Leica Biosystems). Parasagittal serial sections of 6 µm were obtained via a Leica RM2235 rotary microtome. Knee parasagittal sections were stained with hematoxylin and eosin, along with Safranin O and Fast Green (Sigma-Aldrich). Sections were scored by at least two independent observers (Supplementary Table 3), blinded to the experimental groups, as previously described ^12^, using a Leica DM 2000 microscope and Leica Application Suite v4.9 software.

**Supplementary Table 3:** Scoring criteria for histological evaluation of joint pathology.

| Subsynovial Inflammation |  |
| --- | --- |
| 0 | Normal |
| 1 | Focal inflammatory infiltrates, adiposity hardly affected |
| 2 | Focal inflammatory infiltrates equal adiposity |
| 3 | Random inflammatory infiltrates dominating cellular histology |
| 4 | Substantial inflammatory infiltrates with severe loss of adiposity |
| 5 | Ablation of adiposity due to inflammatory infiltrates |
| Synovial Exudate |  |
| 0 | Normal |
| 1 | Evidence of inflammatory cells in space |
| 2 | Moderate numbers of inflammatory cells in space, with evidence of fibrin deposits |
| 3 | Substantial number of inflammatory cells with large fibrin deposits |
| Pannus Formation |  |
| 0 | Normal (between one and three layers) |
| 1 | Over three-layer thick synovial lining and evidence of thickening and/ or invasion of joint space |
| 2 | Over three-layer thick synovial lining 'creeping' over cartilage surfaces and/or finger-like processes into joint space |
| 3 | Over three-layer thick synovial lining showing substantial covering of cartilage surfaces with evident cartilage loss |
| Cartilage and Bone Erosion |  |
| 0 | Normal |
| 1 | Detectable loss of cartilage detected by Safranin O staining |
| 2 | Detectable erosion of underlying bone by pannus activity |
| 3 | Pannus has destroyed a significant part of the bone |

#### LC/MS/MS analysis of oxPLs

Lipid extracts were separated using reverse-phase HPLC on a Luna C_18_ column (150 mm x 2 mm x 3µm) (Phenomenex, Torrance, CA). A gradient elution method of 50 – 100 % B over 10 min followed by 30 min at 100 % B (A, methanol:acetonitrile:water, 1 mM NH_4_CH_3_CO_2_, 60:20:20; B, methanol, 1 mM NH_4_CH_3_CO_2_) was applied with a total flow rate of 200 μl/min. Products were analyzed in multiple reaction monitoring (MRM) mode, on a 6500 Q-Trap (Sciex, Cheshire, United Kingdom), operating in the negative mode, using the following ion source parameters: Temperature: 500 °C, Curtain gas (CUR): 35 psi, Source Gas 1 (GS1): 40 psi, Source Gas 2 (GS2): 30 psi, Ion spray voltage: −4500 V, entrance potential (EP): − 10 V, collision energy (CE) −38 V, declustering energy (DP) −50 V, collision cell exit potential (CXP) −11 V. Transitions were monitored from precursor mass (Q1 *m/z*) to product ion mass (Q3 *m/z*), with a dwell time of 75 msec. For quantification, a mixed isomer HETE-PLs standard curve was generated and a known isomer ratio was used for the determination of each lipid isomer concentration ^3^. MRMs used are provided in Supplementary Table 4. LOD and LOQ were set at 3, and 5:1 respectively (S/N) with at least 6 data points per peak required. Representative chromatograms are shown in Supplementary Figure 15.

#### LC/MS/MS analysis of oxylipins

Lipids were separated using reverse phase HPLC on an Agilent Eclipse Plus C_18_ column (150 mm x 2.1 mm x 1.8 µm) (Phenomenex, Torrance, CA) t 45 °C, with a flow rate of 500 µl/min. A gradient elution method was used where mobile phase B is held at 30 % for 1 minute, then increased to 100 % B from 1 - 17.5 minutes (A: 94.9 % water, 5 % solvent B, 0.1 % glacial acetic acid; B: 84 % acetonitrile, 15.9 % methanol, 0.1 % glacial acetic acid), 100 % B is held from 17.5 - 21 minutes, followed by a decrease to 30 % of B from 21 - 22.5 minutes, which is held until the end of the run at 22.5 minutes. Lipids were analyzed using a scheduled MRM method on a 6500 Q-Trap (Sciex, Cheshire, United Kingdom). A time window is set for the detection of each analyte according to the expected retention time, and transitions are monitored from precursor mass (Q1 *m/z*) to product ion mass (Q3 *m/z*) in negative ion mode, under the following ion source parameters: Temperature: 475 °C, Curtain gas (CUR): 35 psi, Source Gas 1 (GS1): 60 psi, Source Gas 2 (GS2): 60 psi, Ion spray voltage: −4500 V, entrance potential (EP): − 10V. The area under the curve for the precursor ion to product ion transition was integrated using Multiquant 3.0.2. (AB Sciex, Canada) and normalized to the corresponding IS. For quantification, specific isomeric standards were used to generate a standard curve. An equation for calculation was obtained using 1/x^2^ weighted linear regression. MRMs used are provided in Supplementary Table 5. LOD and LOQ were set at 3, and 5:1 respectively (S/N) with at least 6 data points per peak required. Representative chromatograms are shown in Supplementary Figure 16.

#### Chiral LC/MS/MS

Separation was achieved using reversed-phase HPLC on a ChiralPak AD-RH column (150 mm × 4.6 mm x 5 µm; Daicel Corporation) with an isocratic gradient of methanol:water:glacial acetic acid 95:5:0.1 (v/v) with flow rate 300 µl/min for 25 minutes at 40 °C. Products were analyzed in MRM mode, on a 4000 Q-Trap (Sciex, Cheshire, United Kingdom). Transitions were monitored from precursor mass (Q1 *m/z*) to product ion mass (Q3 *m/z*) in negative ion mode, with a dwell time of 125 msec, with the following ion source parameters: Temperature: 500 °C, Curtain gas (CUR): 20 psi, Source Gas 1 (GS1): 40 psi, Source Gas 2 (GS2): 30 psi, Ion spray voltage: −4500 V, entrance potential (EP): − 10V. MRMs used are provided in Supplementary Table 6. LOD and LOQ were set at 3, and 5:1 respectively (S/N) with at least 6 data points per peak required.

#### Statistical analysis

Statistical analysis used Graphpad Prism 9. Shapiro-Wilk test was used for normality testing. Non-parametric analysis was performed using Mann-Whitney or Kruskal–Wallis tests. Statistical differences between conditions were calculated using Dunn’s multiple comparisons test. Where data was normally distributed, Student t-test, one-way or two-way ANOVA was used with Tukey’s Post Hoc test. Heatmaps and hierarchical clustering were used a Pheatmap package in R. Heatmaps display log10 of averaged lipid amounts (ng), normalized to cell count, volume (ml) or wet tissue weight (mg), allowing row-wise and column-wise comparison.

**Supplementary Table 4:**
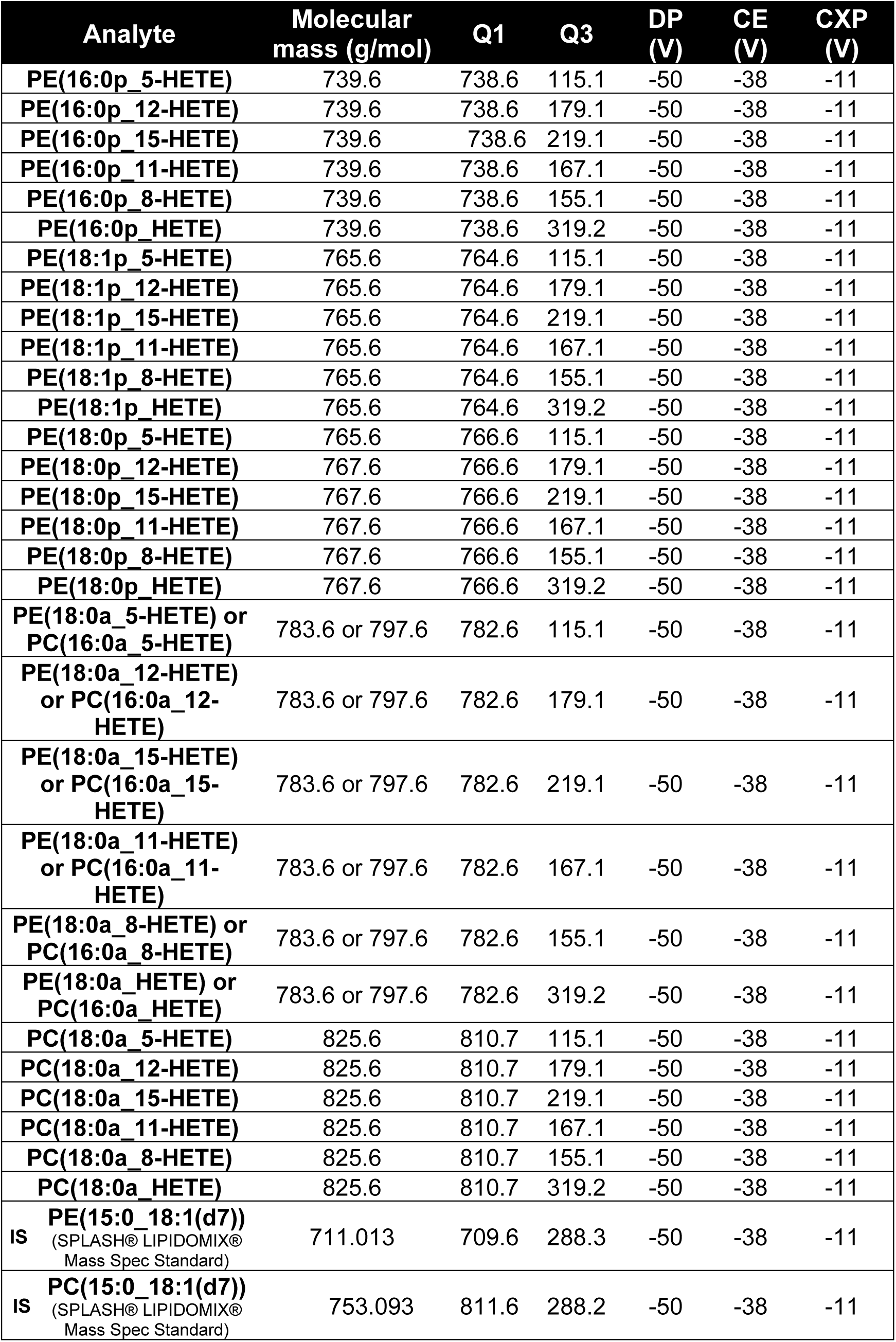
MRM transition for oxPL precursor ion to product ion transitions. Declustering potential (DP), collision potential (CE), Collision cell exit potential (CXP)

| Analyte | Molecular mass (g/mol) | Q1 | Q3 | DP (V) | CE (V) | CXP (V) |
| --- | --- | --- | --- | --- | --- | --- |
| PE(16:0p_5-HETE) | 739.6 | 738.6 | 115.1 | -50 | -38 | -11 |
| PE(16:0p_12-HETE) | 739.6 | 738.6 | 179.1 | -50 | -38 | -11 |
| PE(16:0p_15-HETE) | 739.6 | 738.6 | 219.1 | -50 | -38 | -11 |
| PE(16:0p_11-HETE) | 739.6 | 738.6 | 167.1 | -50 | -38 | -11 |
| PE(16:0p_8-HETE) | 739.6 | 738.6 | 155.1 | -50 | -38 | -11 |
| PE(16:0p_HETE) | 739.6 | 738.6 | 319.2 | -50 | -38 | -11 |
| PE(18:1p_5-HETE) | 765.6 | 764.6 | 115.1 | -50 | -38 | -11 |
| PE(18:1p_12-HETE) | 765.6 | 764.6 | 179.1 | -50 | -38 | -11 |
| PE(18:1p_15-HETE) | 765.6 | 764.6 | 219.1 | -50 | -38 | -11 |
| PE(18:1p_11-HETE) | 765.6 | 764.6 | 167.1 | -50 | -38 | -11 |
| PE(18:1p_8-HETE) | 765.6 | 764.6 | 155.1 | -50 | -38 | -11 |
| PE(18:1p_HETE) | 765.6 | 764.6 | 319.2 | -50 | -38 | -11 |
| PE(18:0p_5-HETE) | 765.6 | 766.6 | 115.1 | -50 | -38 | -11 |
| PE(18:0p_12-HETE) | 767.6 | 766.6 | 179.1 | -50 | -38 | -11 |
| PE(18:0p_15-HETE) | 767.6 | 766.6 | 219.1 | -50 | -38 | -11 |
| PE(18:0p_11-HETE) | 767.6 | 766.6 | 167.1 | -50 | -38 | -11 |
| PE(18:0p_8-HETE) | 767.6 | 766.6 | 155.1 | -50 | -38 | -11 |
| PE(18:0p_HETE) | 767.6 | 766.6 | 319.2 | -50 | -38 | -11 |
| PE(18:0a_5-HETE) or PC(16:0a_5-HETE) | 783.6 or 797.6 | 782.6 | 115.1 | -50 | -38 | -11 |
| PE(18:0a_12-HETE) or PC(16:0a_12-HETE) | 783.6 or 797.6 | 782.6 | 179.1 | -50 | -38 | -11 |
| PE(18:0a_15-HETE) or PC(16:0a_15-HETE) | 783.6 or 797.6 | 782.6 | 219.1 | -50 | -38 | -11 |
| PE(18:0a_11-HETE) or PC(16:0a_11-HETE) | 783.6 or 797.6 | 782.6 | 167.1 | -50 | -38 | -11 |
| PE(18:0a_8-HETE) or PC(16:0a_8-HETE) | 783.6 or 797.6 | 782.6 | 155.1 | -50 | -38 | -11 |
| PE(18:0a_HETE) or PC(16:0a_HETE) | 783.6 or 797.6 | 782.6 | 319.2 | -50 | -38 | -11 |
| PC(18:0a_5-HETE) | 825.6 | 810.7 | 115.1 | -50 | -38 | -11 |
| PC(18:0a_12-HETE) | 825.6 | 810.7 | 179.1 | -50 | -38 | -11 |
| PC(18:0a_15-HETE) | 825.6 | 810.7 | 219.1 | -50 | -38 | -11 |
| PC(18:0a_11-HETE) | 825.6 | 810.7 | 167.1 | -50 | -38 | -11 |
| PC(18:0a_8-HETE) | 825.6 | 810.7 | 155.1 | -50 | -38 | -11 |
| PC(18:0a_HETE) | 825.6 | 810.7 | 319.2 | -50 | -38 | -11 |
| IS PE(15:0_18:1(d7)) (SPLASH® LIPIDOMIX® Mass Spec Standard) | 711.013 | 709.6 | 288.3 | -50 | -38 | -11 |
| IS PC(15:0_18:1(d7)) (SPLASH® LIPIDOMIX® Mass Spec Standard) | 753.093 | 811.6 | 288.2 | -50 | -38 | -11 |

**Supplementary Table 5:** MS/MS parameters for free HETEs: Declustering potential (DP), collision potential (CE), Collision cell exit potential (CXP)

| Analyte | Molecular mass (g/mol) | Q1 | Q3 | Retention time (min) | DP (V) | CE (V) | CXP (V) |
| --- | --- | --- | --- | --- | --- | --- | --- |
| 5-HETE | 320.5 | 319.2 | 115.1 | 14.4 | -55 | -19 | -7 |
| 8-HETE | 320.5 | 319.2 | 155.1 | 14.1 | -65 | -18 | -8 |
| 9-HETE | 320.5 | 319.2 | 167.1 | 14.27 | -50 | -20 | -9 |
| 11-HETE | 320.5 | 319.2 | 167.1 | 13.91 | -60 | -19 | -9 |
| 12-HETE | 320.5 | 319.2 | 179.1 | 14.11 | -65 | -18 | -12 |
| 15-HETE | 320.5 | 319.2 | 219.1 | 13.65 | -55 | -18 | -14 |
| 20-HETE | 320.5 | 319.2 | 275.1 | 12.64 | -85 | -21 | -11 |
| 5-HEPE | 318.4 | 317.2 | 115.1 | 13.17 | -60 | -20 | -10 |
| 8-HEPE | 318.4 | 317.2 | 155.1 | 12.8 | -65 | -19 | -8 |
| 9-HEPE | 318.4 | 317.2 | 167.1 | 12.99 | -50 | -18 | -12 |
| 11-HEPE | 318.4 | 317.2 | 167.1 | 12.69 | -50 | -20 | -13 |
| 12-HEPE | 318.4 | 317.2 | 179.1 | 12.91 | -65 | -18 | -8 |
| 15-HEPE | 318.4 | 317.2 | 219.1 | 12.63 | -65 | -16 | -10 |
| 18-HEPE | 318.4 | 317.2 | 259.1 | 12.25 | -50 | -15 | -11 |
| 4-HDOHE | 344.5 | 343.2 | 101.1 | 14.66 | -50 | -17 | -9 |
| 7-HDOHE | 344.5 | 343.2 | 141.1 | 14.2 | -50 | -21 | -9 |
| 8-HDOHE | 344.5 | 343.2 | 189.1 | 14.31 | -50 | -19 | -9 |
| 10-HDOHE | 344.5 | 343.2 | 153.1 | 13.99 | -55 | -21 | -5 |
| 11-HDOHE | 344.5 | 343.2 | 121.1 | 14.14 | -60 | -18 | -10 |
| 13-HDOHE | 344.5 | 343.2 | 193.1 | 13.87 | -55 | -19 | -9 |
| 14-HDOHE | 344.5 | 343.2 | 205.1 | 13.99 | -45 | -17 | -9 |
| 16-HDOHE | 344.5 | 343.2 | 233.1 | 13.73 | -55 | -17 | -10 |
| 17-HDOHE | 344.5 | 343.2 | 201.1 | 13.79 | -70 | -15 | -10 |
| 20-HDOHE | 344.5 | 343.2 | 241.1 | 13.47 | -55 | -17 | -11 |
| 9-HODE | 296.4 | 295.2 | 171.1 | 13.34 | -85 | -23 | -9 |
| 13-HODE | 296.4 | 295.2 | 195.1 | 13.28 | -85 | -23 | -7 |
| 9-HOTrE | 294.4 | 293.2 | 171.1 | 12 | -60 | -20 | -8 |
| 13-HOTrE | 294.4 | 293.2 | 195.1 | 12.2 | -70 | -22 | -12 |
| 5-HETrE | 322.5 | 321.2 | 115.1 | 15.49 | -70 | -19 | -9 |
| 15-HETrE | 322.5 | 321.2 | 221.1 | 14.29 | -70 | -21 | -11 |
| 9-OxoODE | 294.4 | 293.2 | 185.1 | 14 | -85 | -23 | -13 |
| 13-OxoODE | 294.4 | 293.2 | 195.1 | 13.72 | -85 | -25 | -12 |
| 5-OxoETE | 318.4 | 317.2 | 273.1 | 15.06 | -65 | -20 | -11 |
| 12-OxoETE | 318.4 | 317.2 | 153.1 | 14.36 | -75 | -20 | -10 |
| 15-OxoETE | 318.4 | 317.2 | 113.1 | 14 | -60 | -22 | -8 |
| 9,10-DiHOME | 313.5 | 313.2 | 201.1 | 10.9 | -80 | -29 | -8 |
| 12,13-DiHOME | 313.5 | 313.2 | 183.1 | 10.62 | -80 | -28 | -12 |
| 5,6-DiHETrE | 338.5 | 337.2 | 145.1 | 12.64 | -75 | -24 | -10 |
| 8,9-DiHETrE | 338.5 | 337.2 | 127.1 | 12.14 | -70 | -25 | -8 |
| 11,12-DiHETrE | 338.5 | 337.2 | 167.1 | 11.79 | -65 | -26 | -8 |
| 14,15-DiHETrE | 338.5 | 337.2 | 207.1 | 11.45 | -65 | -25 | -10 |
| 5,6-DiHETE | 336.5 | 335.2 | 115.1 | 11.2 | -60 | -23 | -8 |
| 5,15-DiHETE | 336.5 | 335.2 | 115.1 | 9.92 | -60 | -21 | -9 |
| 8,15-DiHETE | 336.5 | 335.2 | 235.1 | 9.63 | -65 | -22 | -4 |
| 14,15-DiHETE | 336.5 | 335.2 | 207.1 | 10.35 | -65 | -23 | -10 |
| 17,18-DiHETE | 336.5 | 335.2 | 247.1 | 9.97 | -65 | -24 | -8 |
| RvE1 | 350.5 | 349.2 | 195.1 | 3.21 | -65 | -22 | -10 |
| RvD1 | 376.5 | 375.2 | 215.1 | 7.47 | -55 | -23 | -9 |
| RvD2 | 376.5 | 375.2 | 141.1 | 6.8 | -65 | -21 | -11 |
| RvD3 | 376.5 | 375.2 | 147.1 | 6.49 | -65 | -24 | -12 |
| RvD5 | 360.5 | 359.2 | 199.1 | 10.09 | -65 | -22 | -17 |
| LTB3 | 338.5 | 337.2 | 195.1 | 11.5 | -65 | -22 | -8 |
| LTB4 | 336.5 | 335.2 | 195.1 | 10.22 | -70 | -23 | -11 |
| 20-carboxy LTB4 | 366.4 | 365.2 | 347.2 | 3.24 | -80 | -25 | -8 |
| 20-hydroxy LTB4 | 352.5 | 351.2 | 195.1 | 3.55 | -80 | -25 | -8 |
| 6-trans LTB4 | 336.5 | 335.2 | 195.1 | 9.89 | -65 | -23 | -9 |
| LXA4 | 352.5 | 351.2 | 115.1 | 7.32 | -55 | -19 | -10 |
| Mar 1 | 360.5 | 359.2 | 250.1 | 10.1 | -60 | -23 | -11 |
| 7,17-diHDPa | 362.5 | 361.2 | 263.1 | 10.38 | -65 | -20 | -4 |
| 9(10)-EpOME | 296.4 | 295.2 | 171.1 | 14.86 | -80 | -21 | -10 |
| 12(13)-EpOME | 296.4 | 295.2 | 195.1 | 14.74 | -80 | -19 | -8 |
| 5(6)-EET | 320.5 | 319.2 | 191.1 | 15.37 | -60 | -16 | -7 |
| 8(9)-EET | 320.5 | 319.2 | 167.1 | 15.15 | -60 | -15 | -7 |
| 11(12)-EET | 320.5 | 319.2 | 167.1 | 15.15 | -60 | -18 | -8 |
| 14(15)-EET | 320.5 | 319.2 | 219.1 | 14.84 | -65 | -18 | -6 |
| 8(9)-EpETE | 318.4 | 317.2 | 127.1 | 14.2 | -70 | -18 | -8 |
| 11(12)-EpETE | 318.4 | 317.2 | 167.1 | 14.12 | -70 | -15 | -11 |
| 14(15)-EpETE | 318.4 | 317.2 | 207.1 | 14.04 | -70 | -18 | -6 |
| 17(18)-EpETE | 318.4 | 317.2 | 215.1 | 13.7 | -75 | -16 | -10 |
| 7(8)-EpDPA | 344.5 | 343.2 | 113.1 | 15.2 | -60 | -16 | -7 |
| 10(11)-EpDPA | 344.5 | 343.2 | 153.1 | 15.08 | -65 | -15 | -7 |
| 13(14)-EpDPA | 344.5 | 343.2 | 193.1 | 15.02 | -70 | -15 | -7 |
| 16(17)-EpDPA | 344.5 | 343.2 | 233.1 | 14.97 | -55 | -16 | -9 |
| 19(20)-EpDPA | 344.5 | 343.2 | 241.1 | 14.71 | -70 | -18 | -11 |
| PGD1 | 354.5 | 353.2 | 317.2 | 6.65 | -55 | -16 | -8 |
| PGD2 | 352.5 | 351.2 | 271.1 | 6.61 | -50 | -22 | -8 |
| PGD3 | 350.4 | 349.2 | 269.1 | 5.26 | -50 | -17 | -11 |
| PGE1 | 368.5 | 353.2 | 317.2 | 6.53 | -60 | -18 | -10 |
| PGE2 | 352.5 | 351.2 | 271.1 | 6.2 | -60 | -19 | -12 |
| PGE3 | 350.4 | 349.2 | 269.1 | 4.86 | -60 | -17 | -10 |
| PGB2 | 334.4 | 333.2 | 175.1 | 8.82 | -60 | -24 | -10 |
| 13,14-dihydro-15-keto PGE2 | 352.5 | 351.2 | 235.1 | 7.33 | -55 | -19 | -13 |
| 13,14-dihydro-15-keto PGD2 | 352.5 | 351.2 | 207.1 | 8.16 | -50 | -25 | -13 |
| 13,14-dihydro-15-keto PF2α | 354.5 | 353.2 | 113.1 | 7.43 | -55 | -23 | -11 |
| 11β-PGE2 | 352.5 | 351.2 | 271.1 | 6.38 | -55 | -23 | -7 |
| 6-keto PGE1 | 368.5 | 367.2 | 143.1 | 3.22 | -55 | -23 | -9 |
| 8-iso PGE2 | 352.5 | 351.2 | 271.1 | 5.94 | -55 | -21 | -10 |
| 15-deoxy-Δ12,14-PGJ2 | 316.4 | 315.2 | 271.1 | 12.44 | -65 | -18 | -8 |
| 8-iso-15-keto PGF2α | 352.5 | 351.2 | 289.1 | 5.37 | -50 | -23 | -12 |
| PGF2α | 354.5 | 353.2 | 309.2 | 5.89 | -85 | -24 | -9 |
| 6-keto PGF1α | 370.5 | 369.2 | 163.1 | 3.3 | -75 | -26 | -10 |
| Thromboxane B2 | 370.5 | 369.2 | 169.1 | 4.83 | -60 | -22 | -12 |
| IS 11-dehydro Thromboxane B2 | 368.5 | 367.2 | 305.2 | 6.24 | -60 | -20 | -10 |
| IS 13(S)-HODE-d4 | 300.5 | 327.2 | 226.1 | 13.22 | -60 | -25 | -7 |
| IS 5(S)-HETE-d8 | 328.5 | 325.2 | 281.1 | 14.32 | -55 | -19 | -8 |
| IS 12(S)-HETE-d8 | 328.5 | 339.2 | 197.1 | 14.02 | -60 | -20 | -12 |
| IS 15(S)-HETE-d8 | 328.5 | 380.2 | 141.1 | 13.55 | -65 | -22 | -11 |
| IS 20-HETE-d6 | 326.5 | 355.2 | 275.1 | 12.6 | -70 | -21 | -8 |
| IS Leukotriene B4-d4 | 340.5 | 355.2 | 275.1 | 10.17 | -65 | -21 | -9 |
| IS Resolvin D1-d5 | 360.5 | 357.2 | 313.2 | 7.41 | -75 | -18 | -11 |
| IS Prostaglandin E2-d4 | 356.5 | 373.2 | 173.1 | 6.16 | -60 | -23 | -12 |
| IS Prostaglandin D2-d4 | 352.5 | 371.2 | 309.2 | 6.58 | -55 | -23 | -10 |
| IS Prostaglandin F2α-d4 | 358.5 | 331.2 | 167.1 | 5.86 | -80 | -24 | -9 |
| IS Thromboxane B2-d4 | 374.5 | 319.2 | 115.1 | 4.79 | -55 | -22 | -10 |
| IS 11-dehydro Thromboxane B2-d4 | 372.5 | 319.2 | 155.1 | 6.21 | -55 | -21 | -11 |
| IS 11(12)-EET (EpETRe) -d11 | 320.5 | 319.2 | 167.1 | 15.09 | -65 | -18 | -11 |

**Supplementary Table 6:**
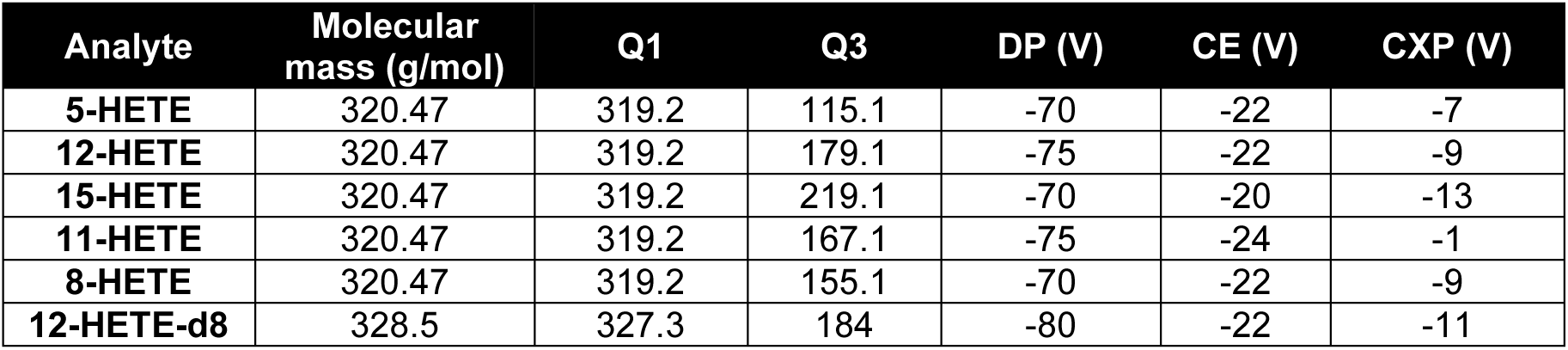
MS/MS parameters for free HETEs for chiral analysis. Declustering potential (DP), collision potential (CE), Collision cell exit potential (CXP)

## Supplementary Data

### Platelet-derived oxylipins are elevated in AIA via IL-6 signaling

Next free oxylipins, including precursors for HETE-PEs, were analyzed. Of 93 measured, only 14 were detected with the most abundant being 12-HETE (Supplementary Figure 7). WT and Il27ra^−/−^mice showed increased 12-HETE on days 3 and 10, while no elevation was observed in Il6ra^−/−^mice (Supplementary Figure 7 B). Other HETEs were present at lower concentrations, with only 11- and 15-HETE quantifiable (Supplementary Figure 7 D,E). 11-HETE displayed a similar pattern to 12-HETE, while 15-HETE peaked on day 10 in Il27ra^−/−^, remaining relatively low in other strains/times (Supplementary Figure 7 D,E). Similar to 12-HETE, other oxylipins, namely 13-HODE (Supplementary Figure 7 C), 14-HDOHE, 9-HODE, 13-HOTrE and 10-HDOHE (Supplementary Figure 7 F-I), peaked at day 3 of AIA in WT mice, displaying a significant increase compared to Il6ra^−/−^. In the case of Il27ra^−/−^ mice, the peak of these oxylipins, with the exception of 9-HODE, occurred on day 10 (Supplementary Figure 7). Leukotrienes, PGs, TXB2, and specialized pro-resolving mediators (SPM) were not reliably detected at either time point, in any strains or in naïve samples. Since many of these can originate from either 12-LOX or COX-1, the data suggest a platelet activation signature in AIA blood, without a PG-associated pro-inflammatory response.

### Generation of HETEs and other oxylipins by blood cells are largely dependent on Alox12

Next, the contribution of LOXs to blood cell oxylipin elevations seen in AIA were determined using Alox15^−/−^ and Alox12^−/−^ blood. Basally, Alox15^−/−^ blood cells contained elevated levels of 12-, 11- and 15-HETEs, as well as 14-HDOHE, 10-HDOHE and 9-HODE, compared to WT (Figure 2 F, Supplementary Figure 8 A). This corresponds with our earlier observation of higher levels of HETE-PEs containing 12-, 15-, and 11-HETEs in Alox15 / blood cells (Figure 2 A-C). Alox12^−/−^ blood cells contained less 12-HETE, as well as other oxylipins basally (Supplementary Figure 8 A). The most striking differences were that following AIA induction, Alox12^−/−^ blood did not show elevations in several oxylipins that were increased in WT blood, including: 12-, 11-HETEs, HDOHEs and HODEs (Supplementary Figures 8), indicating that they were dependent on this isoform. In contrast, the increased levels of 13-HODE in Alox15^−/−^ blood, both basally and during AIA development was somewhat unexpected since 13-HODE can be generated by 12/15-LOX oxidation of linoleic acid (Supplementary Figure 8 G)^15^. This suggests that 13-HODE might be generated either non-enzymatically or by another enzyme following Alox15 deletion. As before, the pattern corresponds with that of eoxPL seen following AIA induction. Although these data show that most oxylipins and eoxPL depend on Alox12^−/−^ in platelets, it’s not always clear which arise directly from enzymatic oxidation versus other mechanisms including (i) indirect non-enzymatic oxidation due to a small amount of radicals exiting the 12-LOX active site during turnover, and (ii) downstream effects such as reduced secondary platelet activation which could impact COX-1. However, the data show that Alox12^−/−^, but not Alox15^−/−^, is required for the oxylipin elevations seen in blood cells in AIA.

**Supplementary Figure 1.**
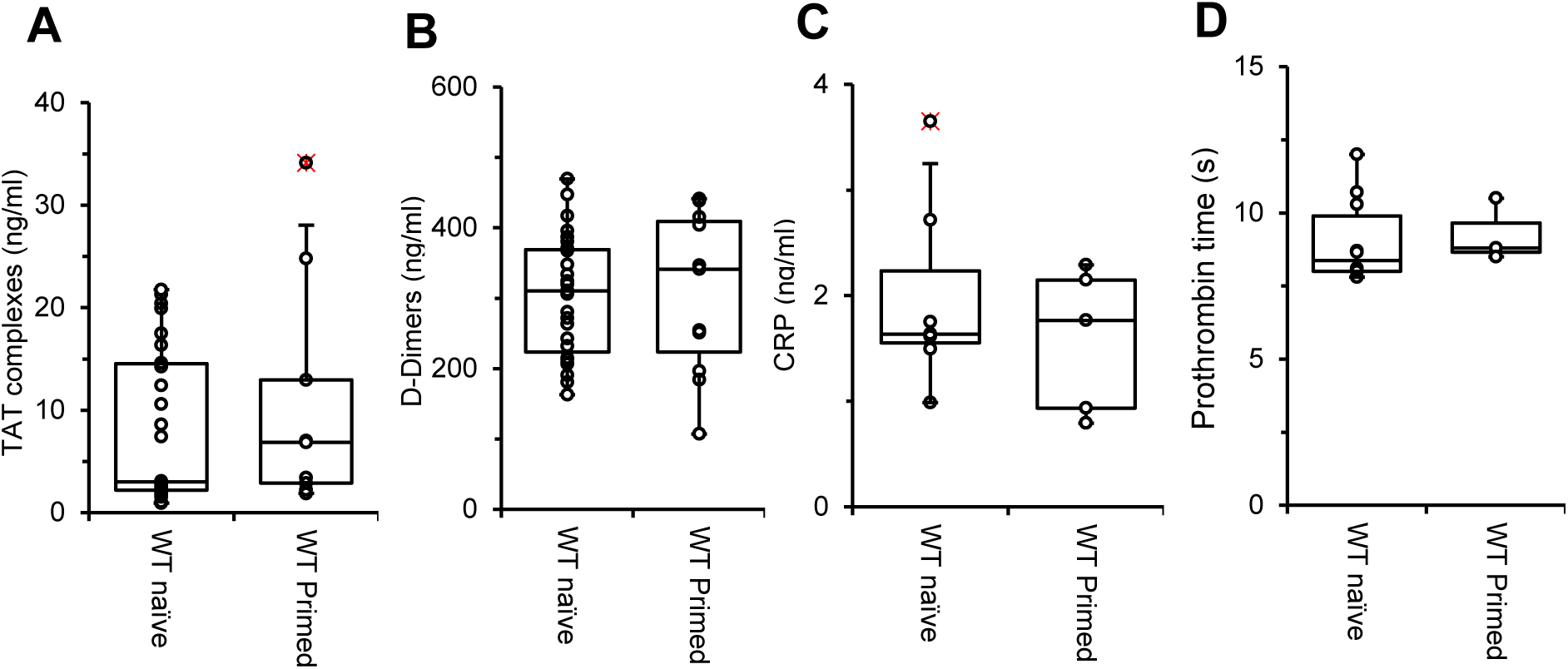
Immunization/priming of wild-type mice does not impact coagulation or inflammation. *WT* mice 8-12 week old were primed using an i.p. injection of Pertussis toxin, and two s.c. injections of mBSA, but induction of arthritis was not performed. *Panel A. Priming does not impact TAT levels.* Plasma levels of TAT complexes were determined, as described in Methods, in WT naïve (n = 30) and primed mice (n = 9). *Panel B. Priming does not impact D-dimer levels.* Plasma levels of D-dimers were determined, as described in Methods, in WT naïve (n = 27) and primed mice (n = 11). *Panel C. Priming does not impact CRP levels.* Plasma levels of CRP were determined as described in Methods in WT naïve (n = 7) and WT primed mice (n = 5). *Panel D. Prothrombin time is not impacted by priming.* Prothrombin time was determined in plasma, as described in Methods, in WT naïve (n = 10) and primed mice (n = 3). Data are represented as box and whisker plots. Data was analyzed using the Mann-Whitney test.

**Supplementary Figure 2.**
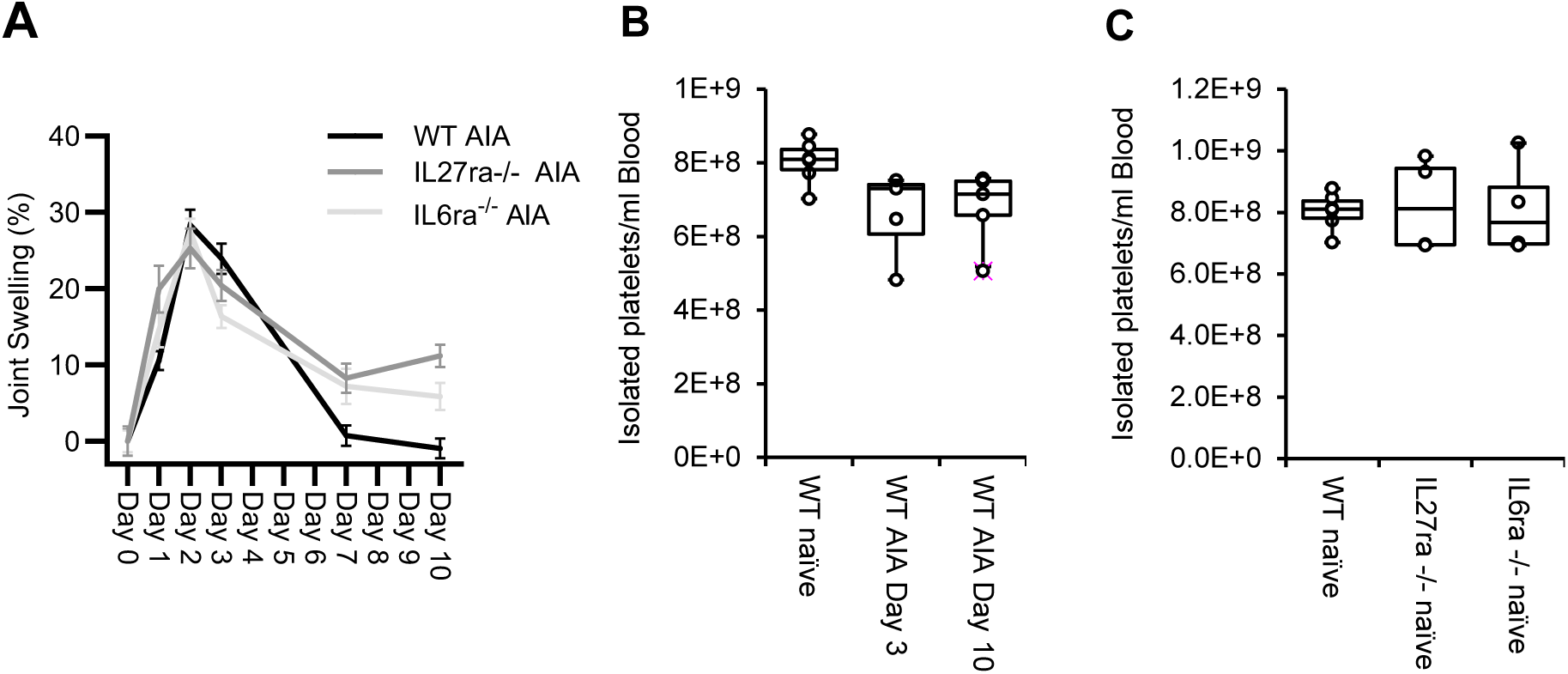
Induction of AIA is not altered in *IL27ra^−/−^* or *IL6ra^−/−^* mice, while platelet count is also unaffected by AIA. *Panel A. Joint swelling confirms the induction of AIA in all strains.* Mice joint diameters of WT (n = 16), IL27ra^−/−^(n = 9) and IL6ra^−/−^(n = 4) mice were measured on day 0, before intra-articular injection, and on days 1, 2, 3, 7 and 10 after mBSA administration. Percentage swelling was calculated with a peak observed between days 2 and 3, confirming the induction of arthritis. *Panel B. AIA model does not significantly alter platelet count.* Platelets from WT mice were isolated and counted at various stages of AIA development. Data are represented as box and whisker plots (n = 4). Data were analyzed using One-way ANOVA and Tukey’s multiple comparison test. *Panel C. Platelet count does not vary between mouse genotypes*. Platelets were isolated and counted from WT (n = 6), IL27ra^−/−^(n = 4), IL6ra^−/−^ (n = 4) mice. Data is represented as box and whisker plots. Data were analyzed using One-way ANOVA and Tukey’s multiple comparison test.

**Supplementary Figure 3.**
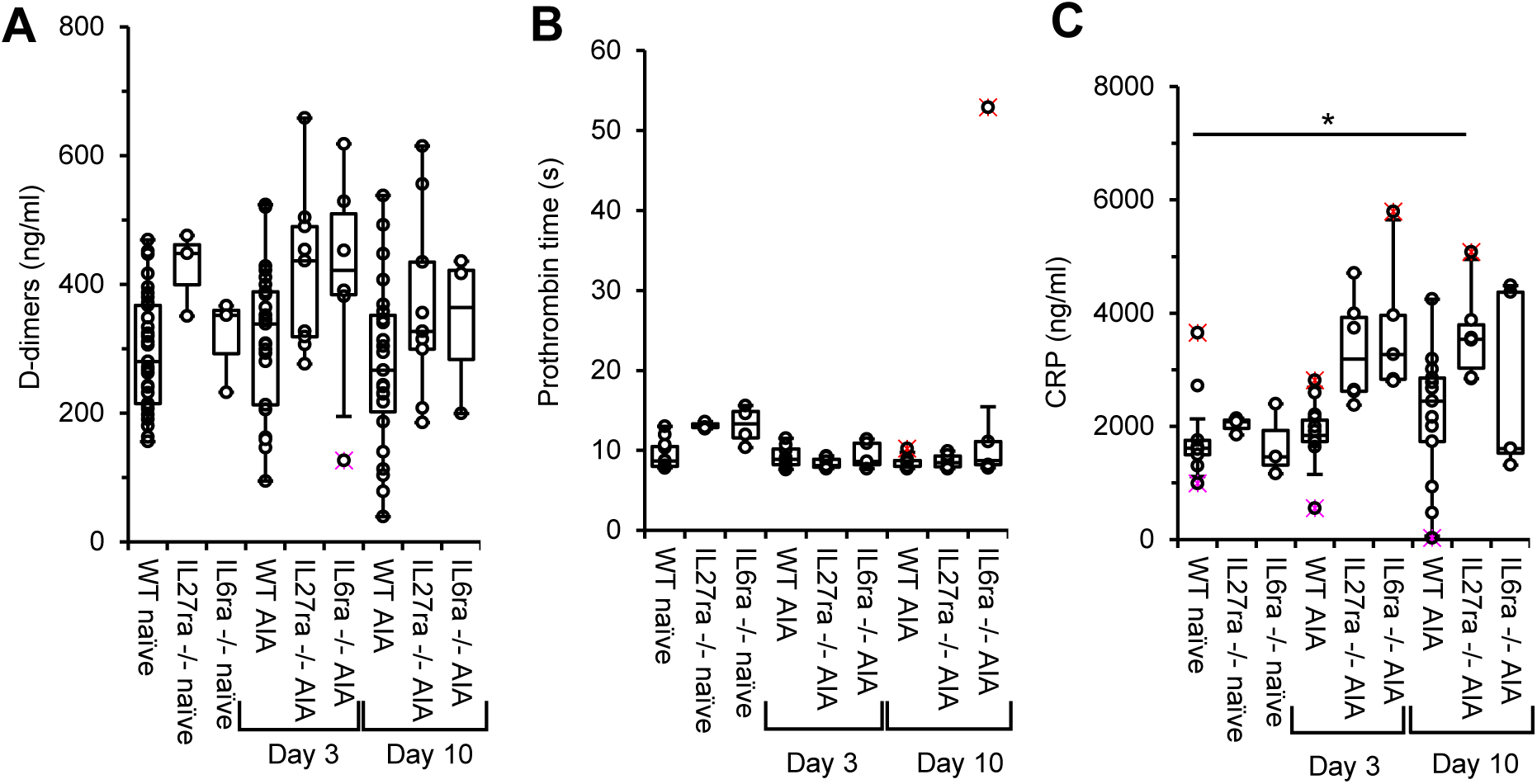
Fibrinolysis, prothrombin time and CRP are not particularly impacted by AIA development. AIA was induced in 8-12 week old *WT*, *Il27ra^−/−^* and *Il6ra^−/−^* male mice. *Panel A. AIA induction in WT, Il27ra^−/−^ and Il6ra^−/−^ does not alter D-Dimer levels.* D-dimers were measured using ELISA. Plasma was collected on day 0 from WT naïve (n = 33), IL27ra^−/−^ naïve (n = 3) and IL6ra^−/−^ naïve (n = 3), as well as on days 3 and 10 of AIA development from WT (n = 25 and 23, respectively), IL27ra^−/−^ (n= 9 for both days) and IL6ra^−/−^ (n= 6 and 4, respectively) mice. *Panel B. AIA induction does not alter prothrombin time.* Prothrombin time was determined as described in Methods. Plasma was collected on day 0 from WT naïve (n = 11), IL27ra^−/−^ naïve (n = 4) and IL6ra^−/−^ naïve (n = 4), as well as on days 3 and 10 of AIA development in WT (n = 8 and 9, respectively), IL27ra^−/−^ (n = 7 and 6, respectively) and IL6ra^−/−^ (n = 5 for both days) mice. *Panel C. CRP is only significant increased on day 10 of AIA development in Il27ra^−/−^ mice.* CRP levels were evaluated by ELISA. Plasma was collected on day 0 from WT naïve (n = 9), IL27ra^−/−^ naïve (n = 3) and IL6ra^−/−^ naïve (n = 3), as well as on days 3 and 10 of AIA development in WT (n = 14 and 13, respectively), IL27ra^−/−^ (n = 6 for both days) and IL6ra^−/−^ (n = 5 for both days) male mice. Data is represented in box and whisker plots. Data were analyzed using the Kruskal-Wallis test and Dunn’s multiple comparison test (* p <0.05, ** p <0.01, *** p<0.001).

**Supplementary Figure 4.**
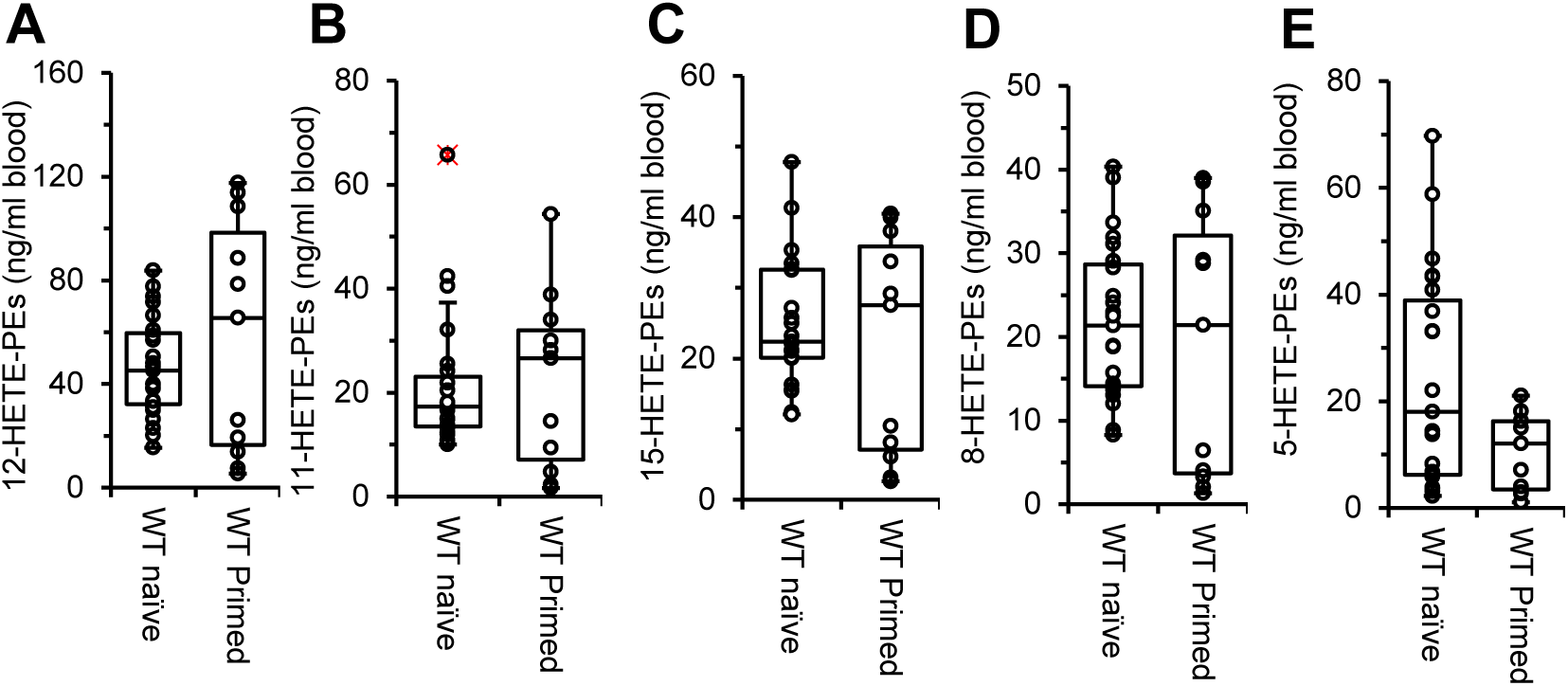
oxPL generation is not significantly altered following immunization/priming alone. *WT* mice, between 8-12 weeks old, were primed using an ip injection of Pertussis toxin, and two s.c. injection of mBSA, without the induction of arthritis. *Panel A-E. 12-, 11-, 15-, 8- and 15-HETE-PEs are not significantly different upon priming.* Whole blood was collected on day 0 from WT naïve (n = 23) and primed (n = 11) mice. Lipids from whole blood cell pellets were extracted as described in Methods and analyzed using LC/MS/MS and the sum of individual HETE-PEs positional isomers was calculated. Data is represented in box and whisker plots. Data were analyzed using Mann-Whitney test.

**Supplementary Figure 5.**
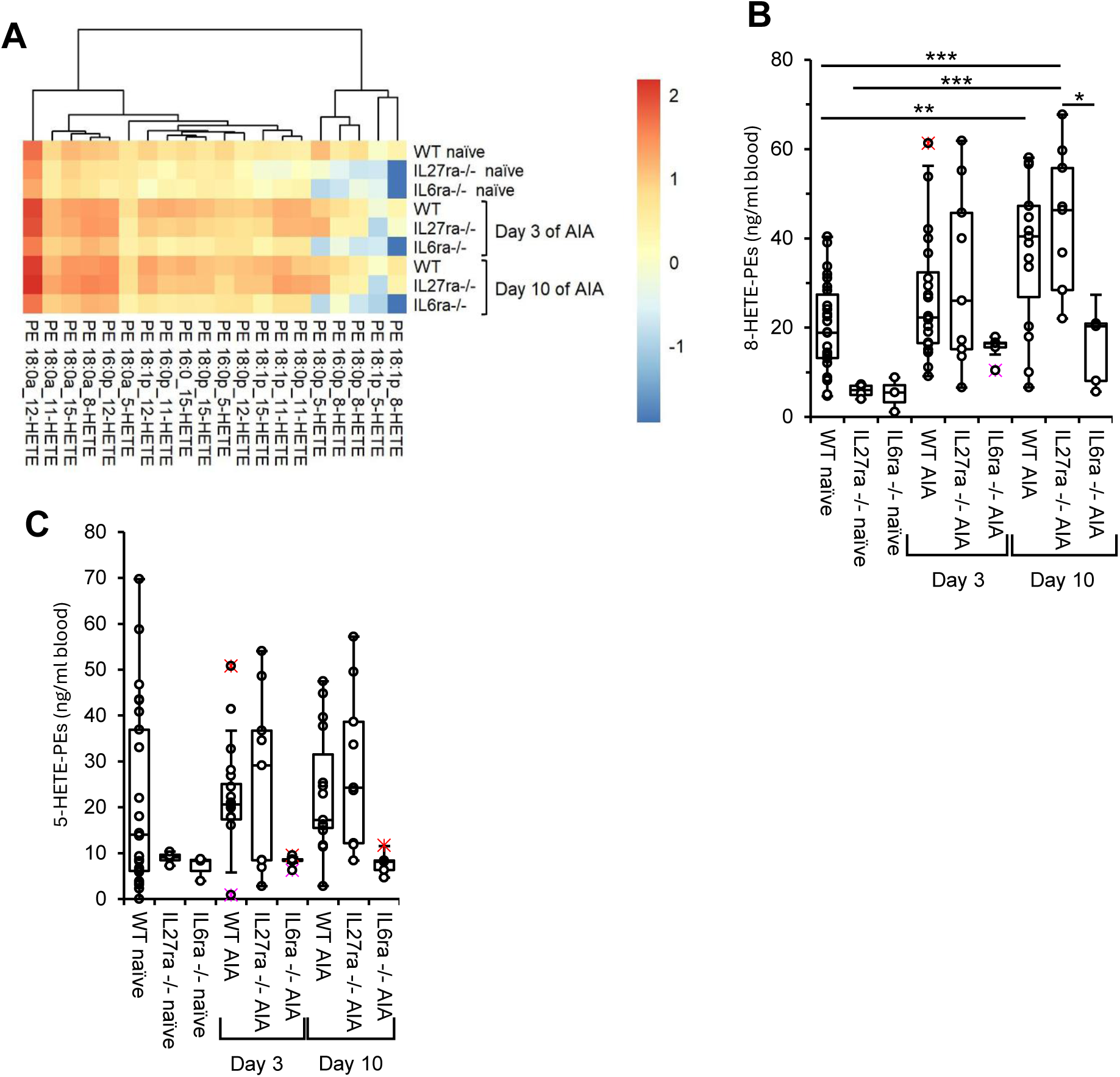
HETE-PEs are elevated in blood cells from WT and *Il27ra^−/−^* during AIA development, especially on day 10. AIA was induced in 8-12 week old *WT, Alox12*^−/−^, *Alox15*^−/−^, *Il27ra*^−/−^ and *Il6ra*^−/−^ male mice as described in Methods, with whole blood collected on Days 3 and 10. Whole blood was collected at day 0 from WT (n = 26), *Il27ra*^−/−^ (n = 4) and *Il6ra*^−/−^ (n = 3), as well as on day 3 and 10 of AIA development from *WT* (n = 23 for day 3; n = 18 for day 10), *Il27ra*^−/−^ (n = 9 for both days) and *Il6ra*^−/−^ (n = 5 for both days) mice. *Panel A. HETE-PEs increase during development of AIA, but not in IL6ra^−/−^ mice.* HETE-PEs were quantified as outlined in Methods using LC/MS/MS. Heatmaps show log10 concentration values (ng/ml) for quantified HETE-PEs. *Panels B,C. 8-HETE-PEs increase during development of AIA, but not in IL6ra^−/−^ mice, while 5-HETE-PEs remain similar throughout.* The sum of 8- and 5-HETE-PEs were quantified as outlined in Methods using LC/MS/MS. Data were analyzed using Two-way ANOVA and Tukey’s multiple comparison test (*p<0.05, **p<0.01, *** p<0.001).

**Supplementary Figure 6.**
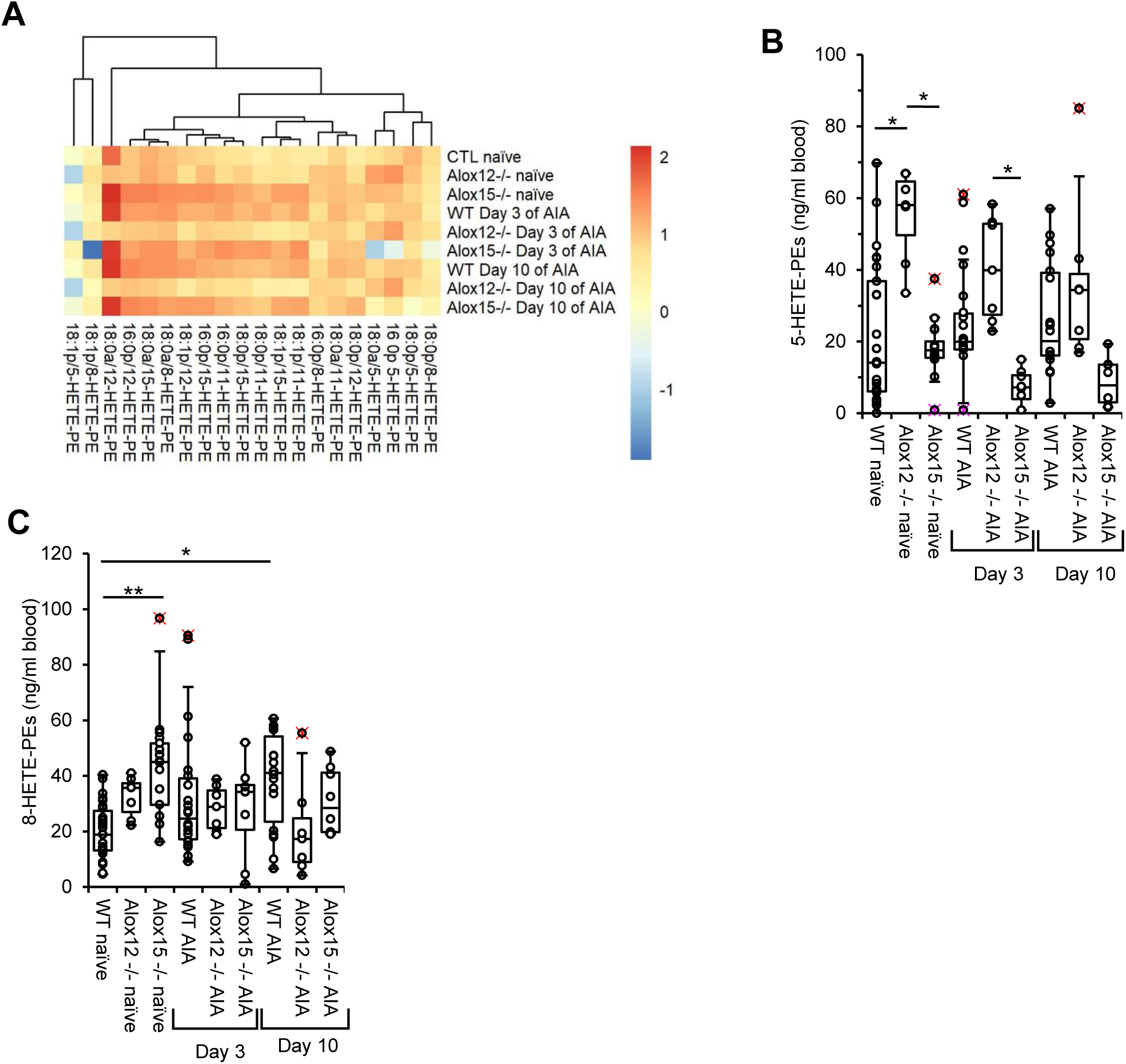
*Alox12,* but not Alox*15* deletion prevents elevation of HETE-PEs levels in mouse blood cells during AIA development. AIA was induced in 8-12 week old male mice as described in Methods, and whole blood collected on day 0 from WT naïve (n = 26), *Alox15*^−/−^ naïve (n = 17) and *Alox12*^−/−^ naïve (n = 7), as well as on day 3 and 10 of AIA from *WT* (n = 23 for day 3; n = 18 for day 10), *Alox15*^−/−^ (n = 8 for both days) and *Alox12*^−/−^ (n = 7 for both days) mice. *Panel A. Alox12 is responsible for the generation of increased levels of eoxPL during AIA.* HETE-PEs were quantified as outlined in Methods using LC/MS/MS. Heatmaps show log10 concentration (ng/ml). *Panels B,C. Alox12 deletion causes basally increased levels of 5-HETE-PE, while Alox15 deletion increased 8-HETE-PEs.* Total 8-HETE-PEs and 5-HETE-PEs were determined as outlined in Methods using LC/MS/MS. Data is represented in box and whisker plots. Data were analyzed using Kruskal-Wallis test and Dunn’s multiple comparisons tests (*p<0.05, **p<0.01, *** p<0.001, **** p<0.0001).

**Supplementary Figure 7.**
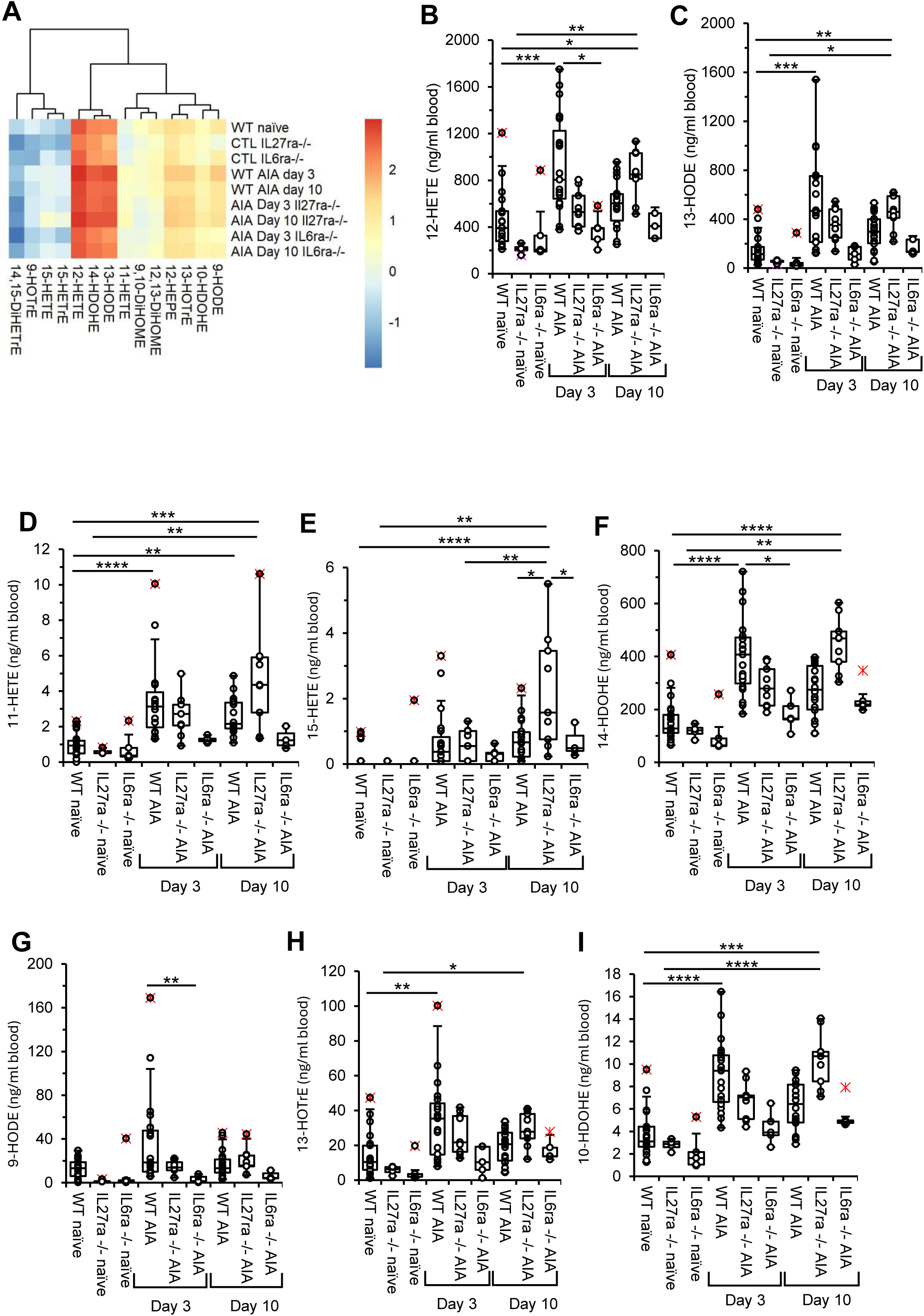
Oxylipins are elevated in blood cells from WT and *Il27ra*^−/−^ during AIA development, especially on day 10. AIA was induced in 8-12 week old *WT, IL27ra*^−/−^ and *IL6ra*^−/−^ male mice as described in Methods. Whole blood was collected at day 0 from WT naïve (n = 21), *IL27ra*^−/−^ naïve (n = 4) and *IL6ra*^−/−^ naïve (n = 5), as well as, on day 3 and 10 of AIA development in *WT* (n = 20 for day 3; n = 15 for day 10), *Il27ra*^−/−^ (n = 9 for both days) and *Il6r*^−/−^ (n = 5 for both days) mice. *Panel A. Oxylipins are increased during AIA development, but not in IL6ra*^−/−^ mice. Oxylipins were quantified as outlined in Methods using LC/MS/MS. Heatmaps show log10 concentration (ng/ml). *Panels B-I. 11- and 15-HETE peak on day 10 of AIA development in IL27ra^−/−^ mice, while 12-HETE, 14-HDODE, 13- and 9-HODE, 13-HOTrE and 10-HDODE peak on day 3 in WT mice.* Lipids from whole blood cell pellets were extracted as described in Methods and analyzed using LC/MS/MS. Total 12-HETE, 13-HODE, 11- and 15-HETEs, 14-HDODE, 9-HODE, 13-HOTrE and 10-HDODE were determined. Data is represented in box and whisker plots. Data were analyzed using Kruskal-Wallis test and Dunn’s multiple comparisons test (* p <0.05, ** p <0.01, *** p<0.001, **** p<0.0001).

**Supplementary Figure 8.**
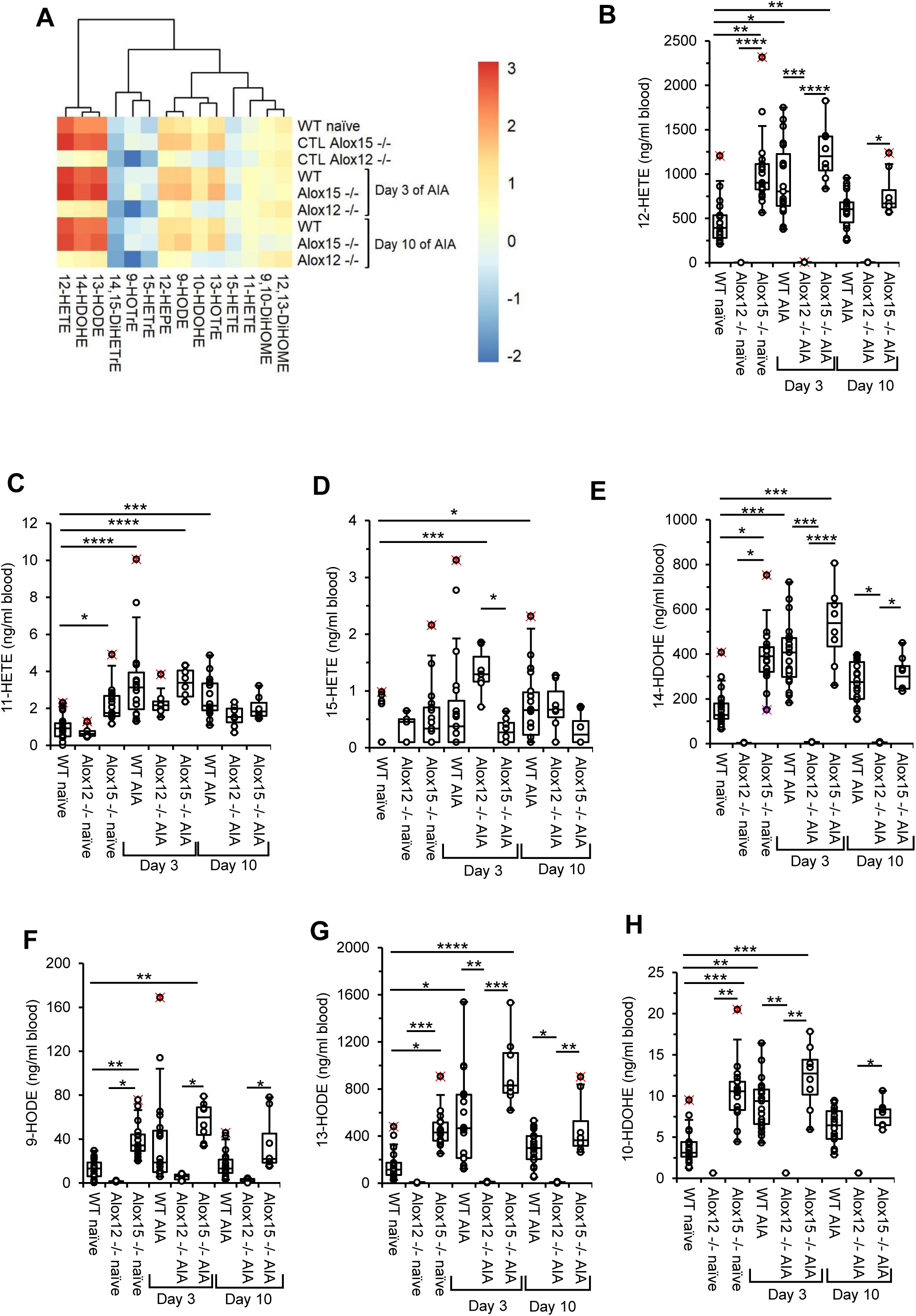
*Alox12* deletion results in the reduction of free HETE*s* and other oxylipins in whole blood during AIA development. AIA was induced in 8-12-week old *WT, Alox12*^−/−^ and *Alox15*^−/−^ male mice as described in Methods. Whole blood was collected at day 0 from WT naïve (n = 26), *Alox15* ^−/−^ naïve (n = 17) and *Alox12*^−/−^ naïve (n = 7), as well as, on day 3 and 10 of AIA development in *WT* (n = 23 for day 3; n = 18 for day 10), *Alox15*^−/−^ (n = 8 for both days) and *Alox12*^−/−^ (n = 7 for both days) mice. *Panel A. Alox12 is responsible for oxylipin generation in blood cells during AIA.* Oxylipins were quantified as outlined in Methods using LC/MS/MS. Heatmaps show log10 concentration values (ng/ml). *Panels B-F. Alox12 deletion decreases generation of 12-HETE, 14-HDODE, 9-HODE and 10-HDODE, while increasing 15-HETE during AIA development.* Lipids from whole blood cell pellets were extracted as described in Methods and analyzed using LC/MS/MS. 11- and 15-HETE, 14-HDODE, 9- and 13-HODEs and 10-HDODE were determined. Data is represented in box and whisker plots. Data were analyzed using Kruskal-Wallis test and Dunn’s multiple comparisons test (* p <0.05, ** p <0.01, *** p<0.001, **** p<0.0001).

**Supplementary Figure 9.**
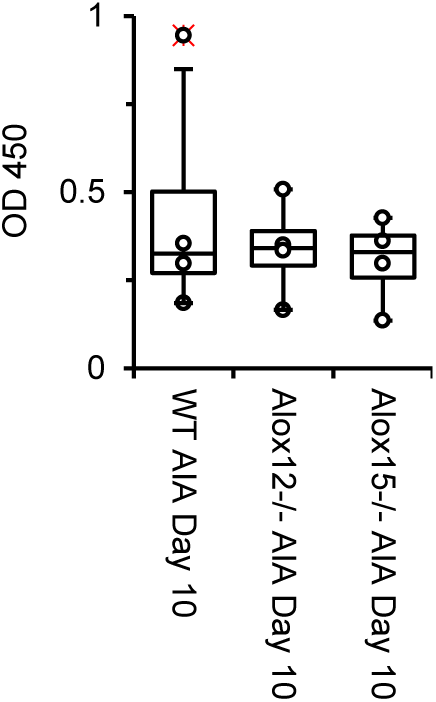
Antibody response to mBSA is similar for WT, *Alox15^−/−^* mice and *Alox12^−/−^* mice. Specific antibody titres against mBSA were determined using ELISA in WT and *Alox15*^−/−^ mice plasma on day 10 post arthritis induction. Data represents mean ± SEM (n = 4) and statistical analysis was performed using a student t-test.

**Supplementary Figure 10.**
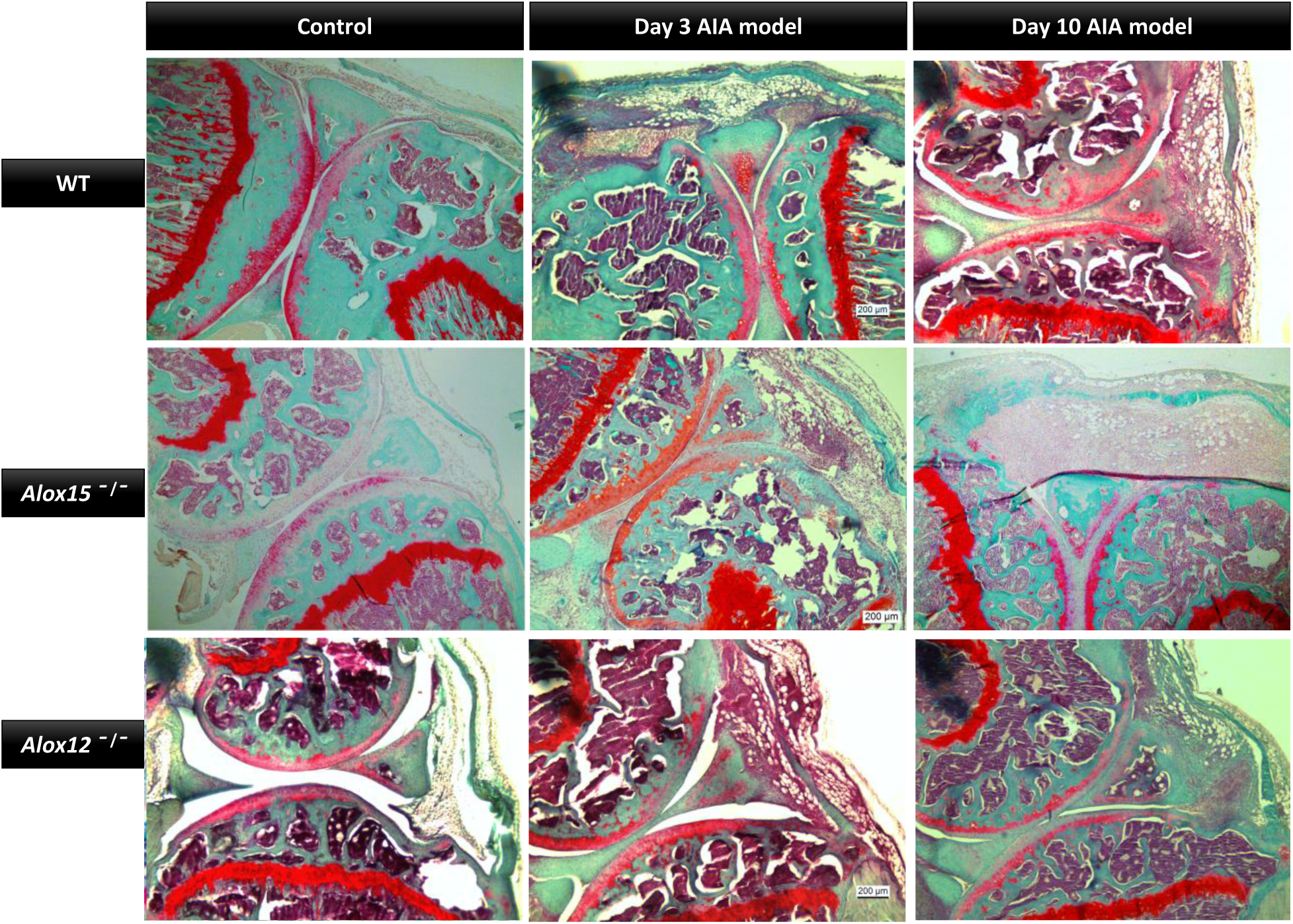
AIA development in both *Alox15^−/−^ and Alox12^−/−^* mice results in a worse phenotype. AIA was induced in 9-12-week old *WT, Alox12*^−/−^ and *Alox15*^−/−^ male mice as described in Methods, with synovial tissue collected on Days 3 and 10. Knee joints were also collected on day 0 from WT, *Alox12*^−/−^ and *Alox15*^−/−^ mice for histological staining and assessment, as described in Methods. Representative images of haematoxylin, fast green and safranin O staining of WT (top), *Alox12^−/−^* (middle) and *Alox15^−/−^* (bottom) mouse knee joints as controls (left), and at days 3 (centre) and 10 (right) of AIA development.

**Supplementary Figure 11.**
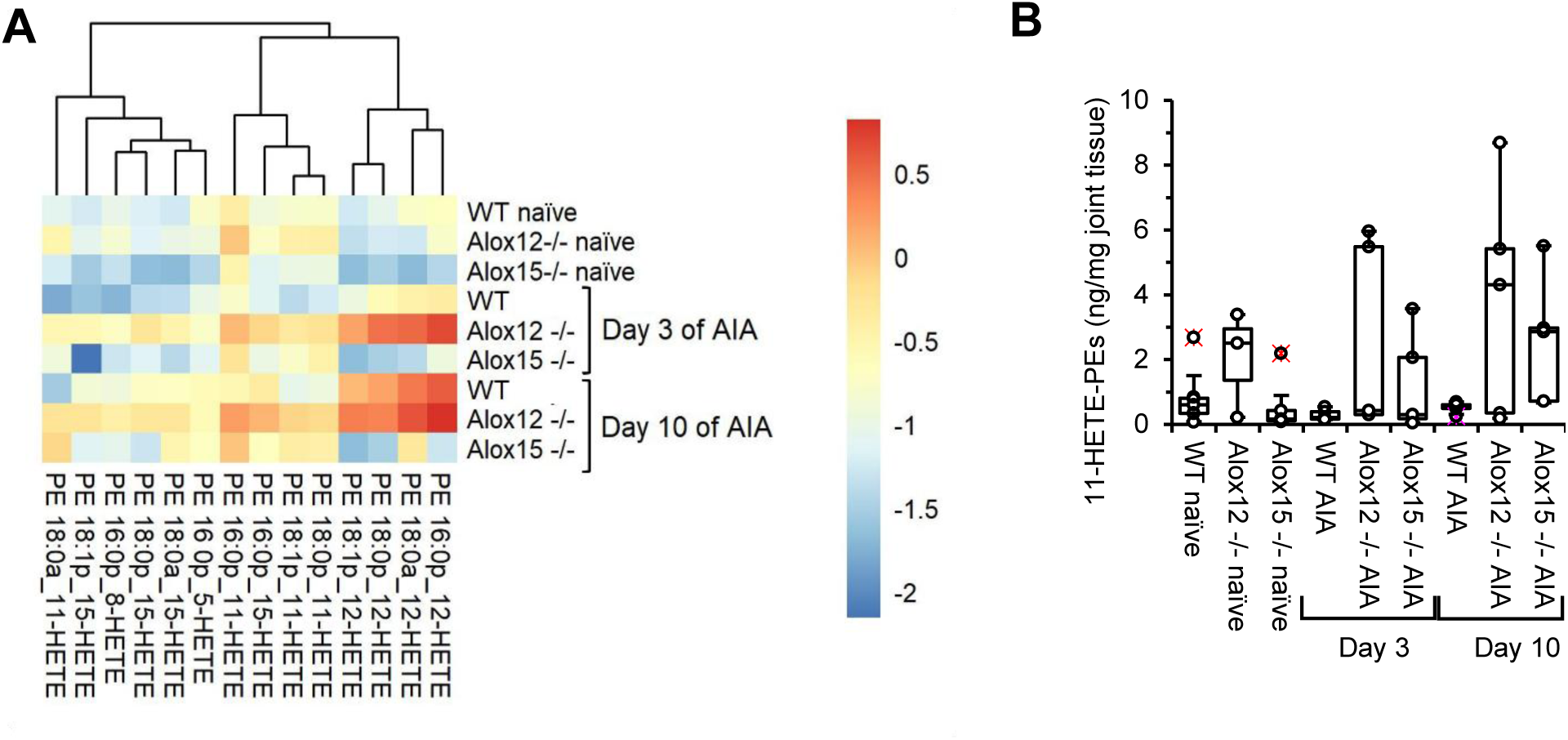
*Alox15* deletion results in *an overall* reduction of oxPLs in synovial tissue during AIA development. AIA was induced in 9-12 week old WT, *Alox12^−/−^* and *Alox15^−/−^* male mice as described in Methods, with knee joints collected on day 0 from WT naïve (n = 7), *Alox15*^−/−^ naïve (n = 5) and *Alox12*^−/−^ naïve (n = 3), as well as, on day 3 and 10 of AIA development from *WT* (n = 5), *Alox15*^−/−^ (n = 5) and *Alox12*^−/−^ (n = 5) mice. Lipids from pooled synovial tissue were extracted as described in Methods and analyzed using LC/MS/MS. *Panel A. Alox15 is responsible for the generation of increased 12- and 15-HETE-PE species during AIA development in synovial tissue.* Heatmaps shows log10 concentration values [ng/mg (wet tissue)] for HETE-PEs species. *Panel B. Deletion of Alox15 or Alox12 does not alter the generation of 11-HETE-PEs during AIA development in synovial tissue.* The sum of 11-HETE-PEs was quantified as outlined in Methods using LC/MS/MS. Data is represented in box and whisker plot. Data were analyzed using Kruskal-Wallis test.

**Supplementary Figure 12.**
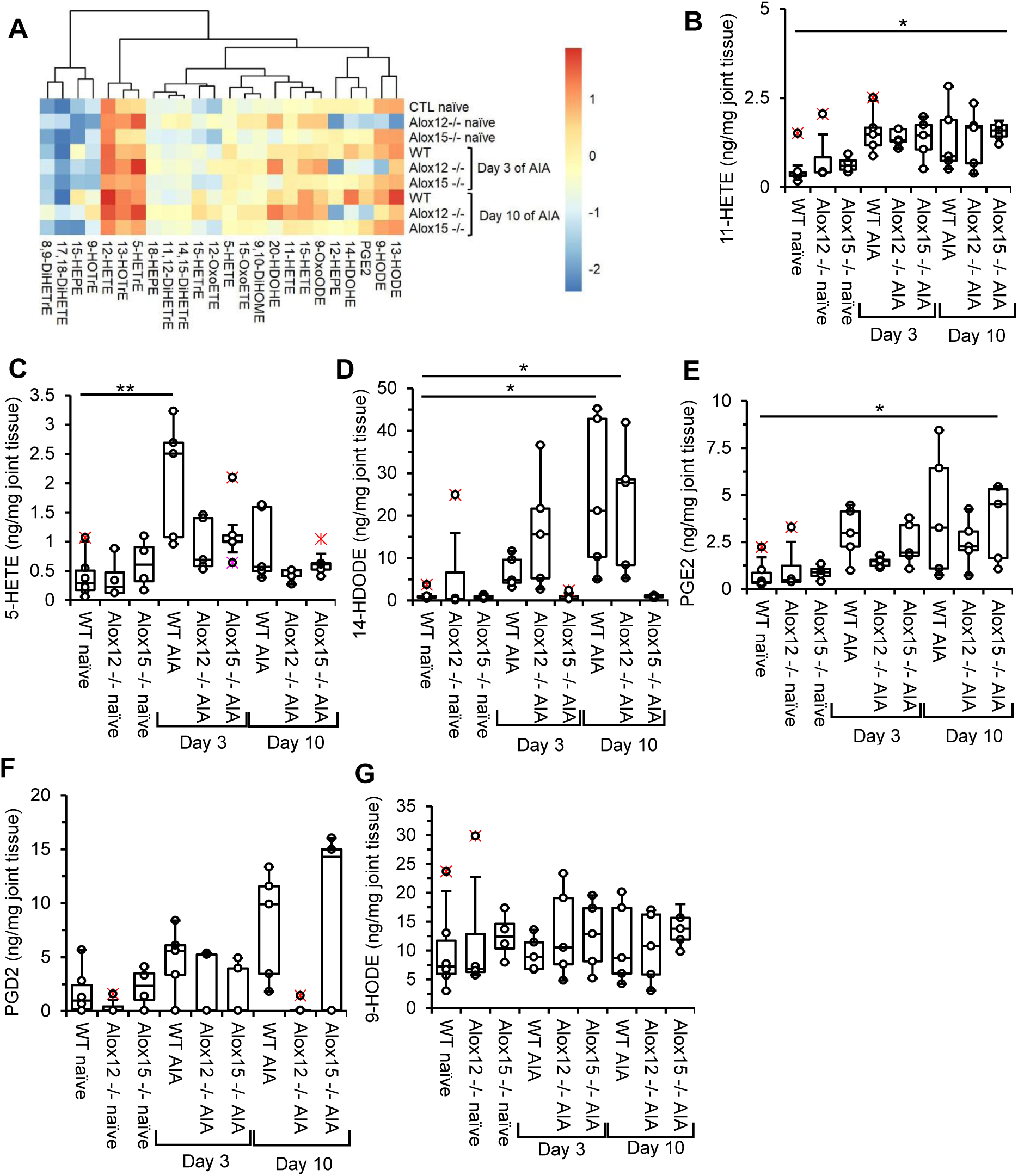
*Alox15* deletion results in *a* reduction of oxylipins in synovial tissue during AIA development. AIA was induced in 9-12 week old WT, *Alox12^−/−^* and *Alox15^−/−^* male mice as described in Methods, with knee joints collected on day 0 for WT (n = 7), *Alox15*^−/−^ (n = 5) and *Alox12*^−/−^ (n = 3), as well as, on day 3 and 10 of AIA development in *WT* (n = 5), *Alox15*^−/−^ (n = 5) and *Alox12*^−/−^ (n = 5) mice. Lipids from pooled synovial tissue were extracted as described in Methods and analyzed using LC/MS/MS. *Panel A. Oxylipin profiles during AIA development is altered by LOX deletion.* Heatmap shows log10 concentration values [ng/mg (wet tissue)]. *Panel B-G. Deletion of Alox15 significantly alters the production of oxylipins during AIA development.* The concentration of oxylipins in synovial tissue (ng/mg tissue) were determined as described in Methods and analyzed using LC/MS/MS, namely 11-HETE and 5-HETE, 14-HDODE, PGE2, PGD2 and 9-HODE. Data is represented in a box and whisker plot. Data were analyzed using Kruskal-Wallis’s test and Dunn’s multiple comparisons test (* p <0.05, ** p <0.01).

**Supplementary Figure 13.**
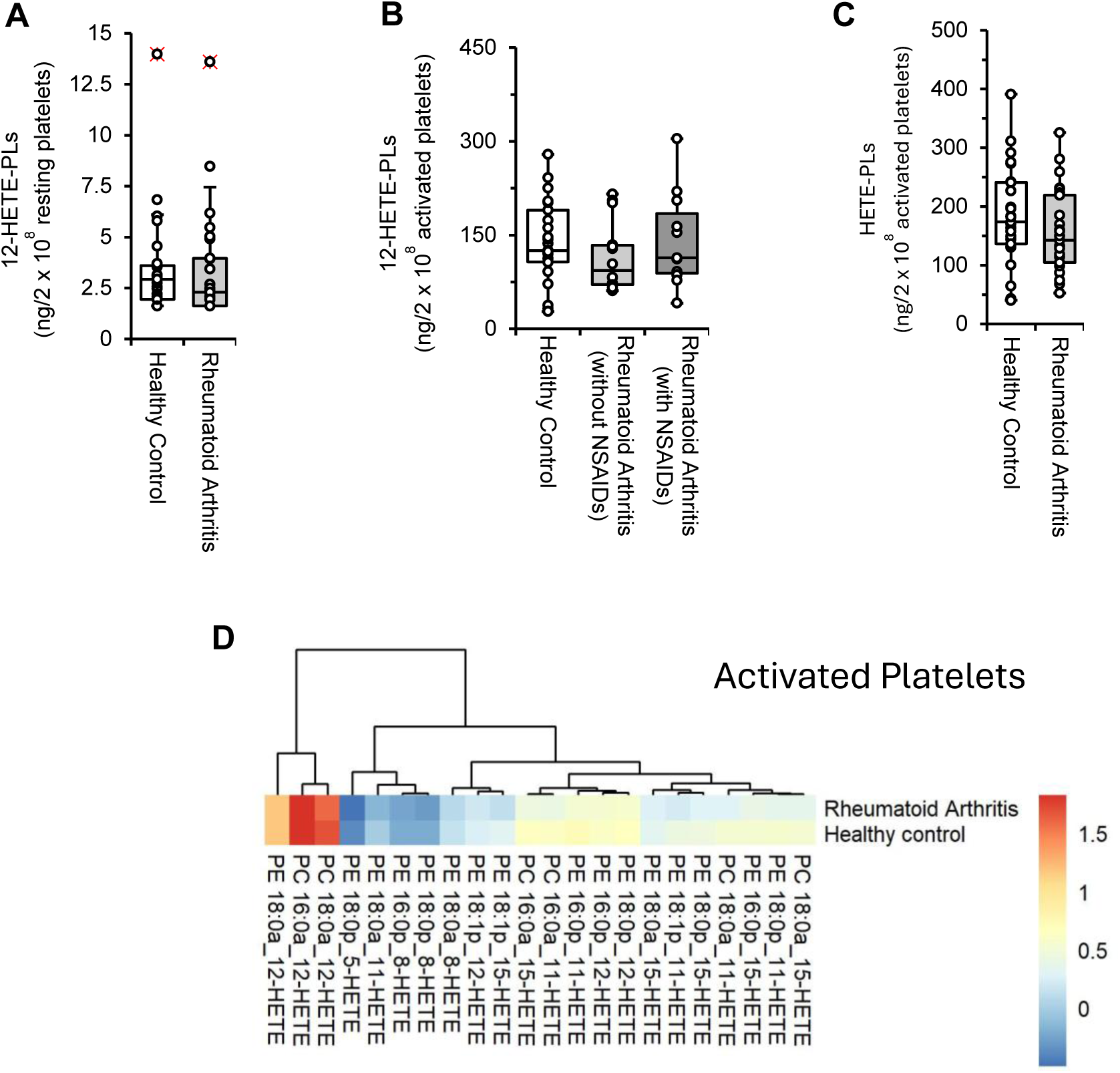
HETE-PLs are similar in HC and RA, and not significantly impacted by aspirin. Washed platelets isolated from RA patient (n=25) and healthy control (n=25) blood were analyzed for eoxPLs using LC/MS/MS. *Panel A. 12-HETE-PL levels in resting platelets are similar between RA and HC.* The sum of 12-HETE-PLs isomers was calculated for RA and HC platelets. Data were analyzed using student’s t-test. *Panel B. NSAID intake does not significantly alter 12-HETE-PLs in activated platelets.* Washed platelets from RA patients taking NSAIDs (n = 11), or not (n = 14), along with HC (n = 25) were isolated and activated using 0.2 U/ml thrombin and 1 mM CaCl_2_, as described in Methods. 12-HETE-PLs were analyzed using LC/MS/MS. Data were analyzed using the Kruskal-Wallis test. *Panel C. Total amount of HETE-PLs in activated platelets are similar between RA and HC.* Platelets were activated with 0.2 U/ml thrombin and 1 mM CaCl_2_. OxPL were analyzed using LC/MS/MS. The sum of all HETE-PLs isomers was calculated for RA and HC platelets. Data were analyzed using Student’s t-test. *Panel C. HETE-PL profile in activated platelets appears reduced in RA patients compared to healthy controls.* Heatmap shows log10 values for analyte amount (ng/2×10^8^ platelets).

**Supplementary Figure 14.**
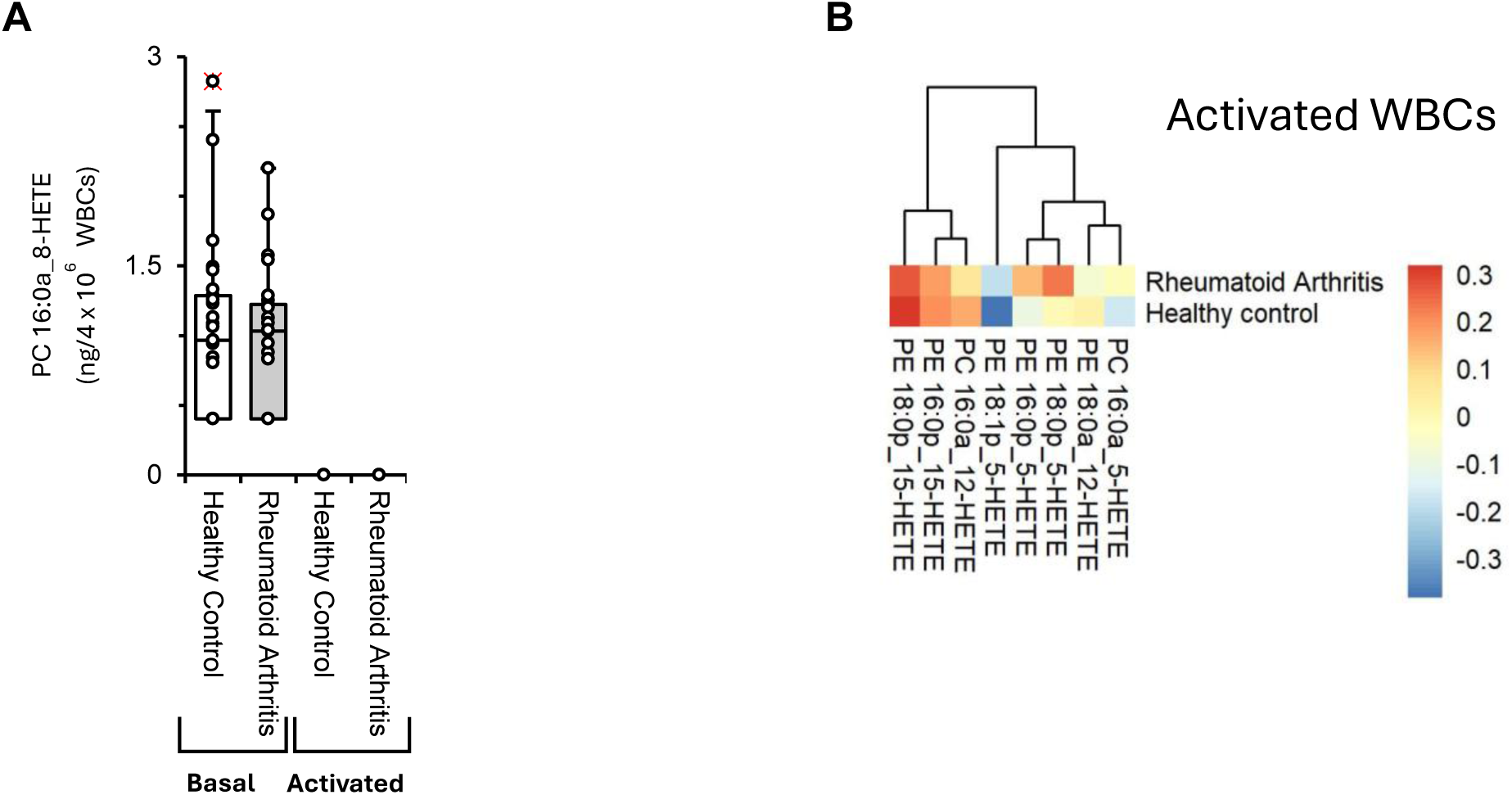
Ionophore activates WBC HETE-PL generation, but RA and HC levels are similar. WBC were isolated and counted as described in Methods, from RA patients and healthy controls from the clinical cohort were analyzed by LC/MS/MS. Resting WBCs from RA patients (n = 25) and healthy controls (n = 25) were isolated as described in Methods, and oxPLs species were analyzed by LC/MS/MS. Isolated WBCs were activated with 10 μM Ca^2+^ Ionophore A23187 and 1 mM CaCl_2_. OxPLs of ionophore activated WBCs from RA patients (n = 25) and healthy controls (n = 25) were analyzed by LC/MS/MS as described in Methods. *Panel A. PC 16:0a_8-HETE is the only HETE-PLs specie detected in resting WBCs from both RA patients and healthy control in similar levels.* The amount of *PC 16:0a_8-HETE* was calculated (ng/4×10^6^ WBCs) in both basal and activated state, in WBCs from RA and healthy controls. *Panel B. Limited species of HETE-PLs were found in activated WBCs in RA patients and healthy control.* Heatmap shows log10 values for analyte amount (ng/4×10^6^ WBCs). Data were analyzed using Unpair t-test between basal WBCs of RA and healthy controls.

**Supplementary Figure 15.**
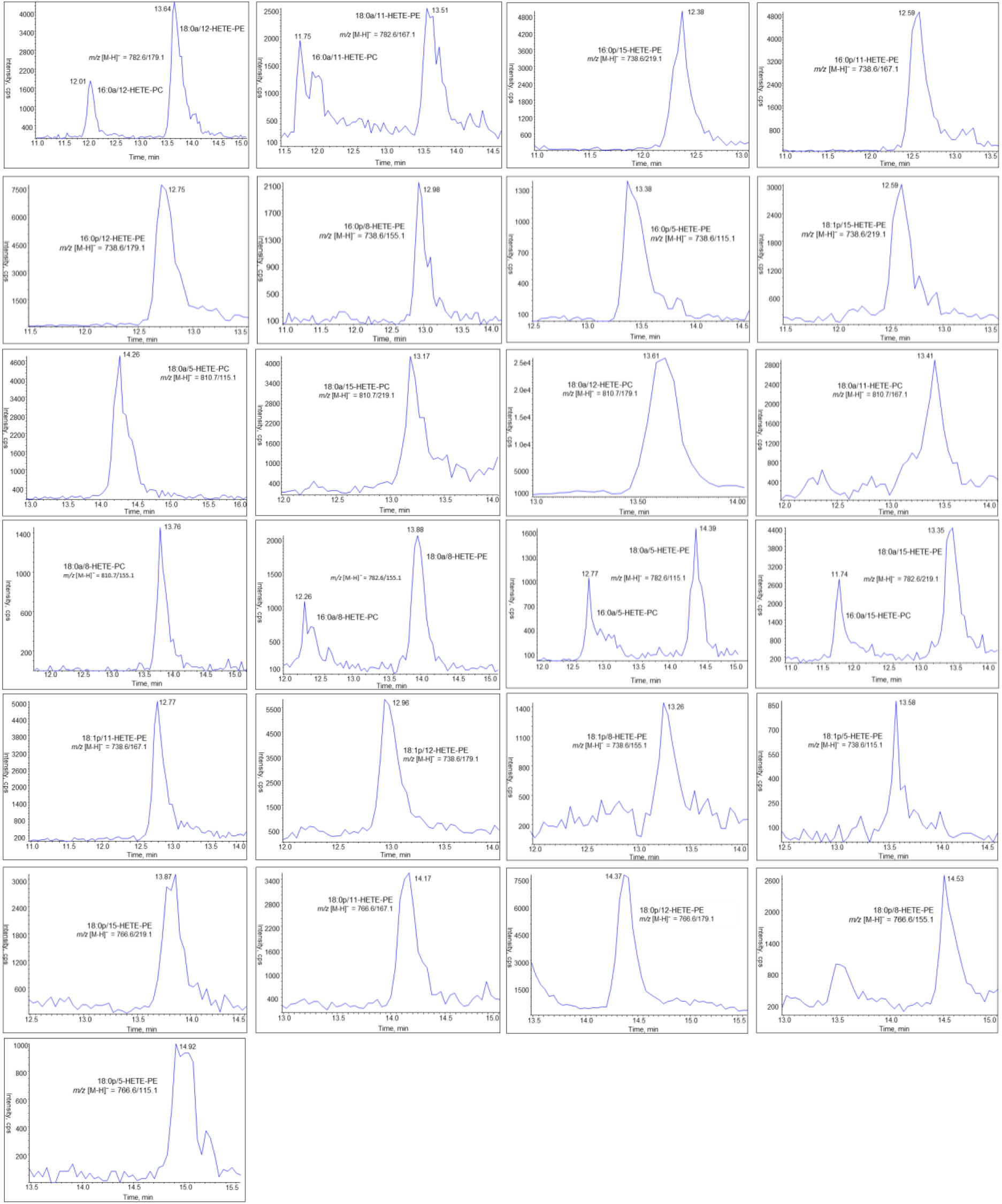
Representative chromatograms of the LC-MS/MS analysis of oxidised phospholipids. Lipid extracts were separated using reverse-phase LC/MS/MS, as described in Methods. Screenshots were taken from Multiquant software.

**Supplementary Figure 16.**
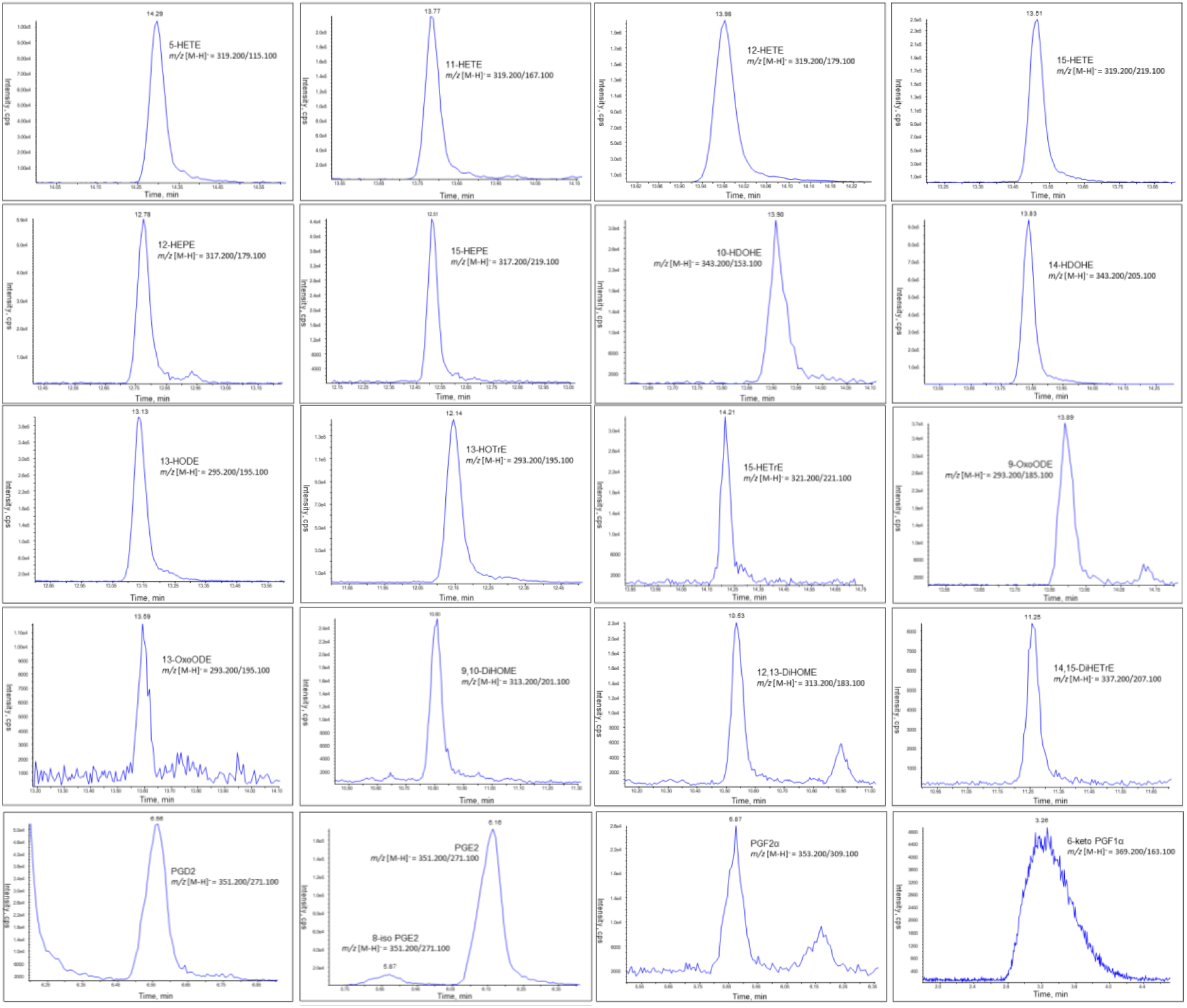
Representative chromatograms of oxylipin LC/MS/MS. Lipid extracts were separated using reverse-phase LC/MS/MS, as described in Methods. Screenshots were taken from Multiquant software. Lipids were confirmed by comparing retention time with primary standards run in the same analytical batch.

